# Large extracellular vesicles regulate endothelial angiogenic potential via paracrine and autocrine signaling

**DOI:** 10.64898/2025.12.17.694905

**Authors:** Grace Richmond, Rose Nguyen, Alanna Sedgwick, Jeffrey S. Schorey, Crislyn D’Souza-Schorey

**Affiliations:** Department of Biological Sciences, University of Notre Dame, Notre Dame, IN 46556

**Keywords:** cancer, endothelial cells, melanoma, extracellular vesicles, tumor microenvironment

## Abstract

Angiogenesis, a process typically associated with tumor growth and development, is often linked to advanced disease and poor clinical outcomes. Tumor cells establish a pro-angiogenic microenvironment through the release of paracrine signaling mediators including extracellular vesicles (EVs). EVs have been shown to facilitate intercellular communication and encompass a diverse range of secreted vesicles, including small EVs (sEVs) which range in size from ∼60 to 100nm and large EVs (L-EVs) which are even more diverse and range from 200nm to >1μm in size. Despite advancements in anti-angiogenic cancer therapies, such as bevacizumab, late-stage tumors, including advanced melanomas, exhibit mixed clinical responses. In this study, we elucidate a unique role for melanoma-derived L-EVs in promoting bevacizumab-insensitive endothelial angiogenic phenotypes. This L-EV-mediated increase in endothelial tube formation is sensitive to the effects of sorafenib, a multi-kinase inhibitor, but not SU5416, a selective VEGF-receptor inhibitor. We also demonstrate that melanoma L-EVs contain VEGF as luminal cargo and induce paracrine effects by modulating the endothelial EV secretome. The release from endothelial cells of soluble VEGF, EVs, and pro-angiogenic cytokines such as IL-8, MIF, and PAI-1 drive sustained endothelial tube formation through autocrine signaling. Finally, we show that EV subtypes have distinct effects on the acquisition of angiogenic phenotypes and their roles vary with tumor type. These findings provide new insight into the mechanisms of angiogenic therapy resistance in melanoma and demonstrate the differential functions of EV subtypes in angiogenesis across tumor types.

## INTRODUCTION

Melanoma is the deadliest form of skin cancer with high rates of metastasis and rising incidence over the last decade. Primary melanoma tumors originate from the transformation of melanocytes in association with UV exposure and grow vertically over time (1–3). Following vertical growth into dermal skin layers, melanoma tumors invade outward and metastasize with support from blood vessel accumulation into dense vascular networks. Melanoma tumor vessel density has a strong clinical correlation to poor prognosis and outcomes of decreased overall survival and high rates of recurrence (4, 5).

Tumor cells drive angiogenesis through paracrine signaling by releasing signaling molecules including VEGF, IL-6, and IL-8 that help coordinate the growth and organization of new blood vessels (6, 7). These pro-angiogenic mediators are released from tumor cells as both soluble form and in extracellular vesicles. Extracellular vesicles (EVs) released from tumor cells can interact with nearby cells to deliver protein, lipid, and nucleic acid cargo that may influence the behavior of recipient cells (8–10). Tumor-EV signaling leads to the conditioning of the tumor microenvironment (TME) in part by facilitating cancer-associated fibroblast reprogramming and modulating the activities of immune cells and endothelial cells (11–14). Studies on angiogenesis have implicated EV-associated factors in driving angiogenic phenotypes, but the mechanisms and responsible cargos are not universal across tumor types. Understanding differences in tumor-derived EV contributions is a complex and lingering gap in the field. Most studies have primarily focused on defining the roles of small EVs in angiogenesis. Small EVs, including exosomes, are a subtype of EV which belongs to a larger family of vesicles that also includes large EVs such as microvesicles (MVs) (15, 16). The roles of the diverse EV subtypes in angiogenesis are relatively underexplored and less understood. Elucidating the roles of EVs in the TME on the modulation of endothelial cell behaviors is important to more fully understand the underlying mechanisms and regulation of tumor angiogenesis.

The importance of understanding the role of EVs is underscored by their influence in clinical therapies targeting tumor angiogenesis, particularly those that have had minimal success and often result in drug resistance and tumor recurrence (17, 18). For instance, reports suggest that treatment strategies targeting VEGF with monoclonal antibodies to block angiogenesis are compromised by the EV-mediated transfer of therapy-insensitive VEGF to neighboring cells (19, 20). Small EV-associated VEGF isoforms were still able to bind receptors on endothelial cells and stimulate pro-angiogenic phenotypes in the presence of bevacizumab. Understanding the impacts of EV subtypes on angiogenesis and how they may influence targeted therapies will be highly useful to inform the development of more effective treatments. Here, we describe the role for melanoma-derived L-EVs in the induction of endothelial angiogenic phenotypes. We have compared the effects of small EVs (sEVs) and large EVs (L-EVs) derived from melanoma and other tumor cell lines on endothelial cells. We show that L-EVs promote endothelial tube formation via mechanisms distinct from sEVs derived from the same cell lines. Furthermore, we demonstrate that the roles of EV subtypes vary with tumor cell type. Immunoassays and enzymatic digestion experiments reveal that melanoma L-EVs contain VEGF as a luminal cargo and promote endothelial tube formation in a bevacizumab-insensitive manner. Additionally, L-EVs promote the release of pro-angiogenic cytokines and EVs from endothelial cells. Further, cytokine neutralization experiments validate that changes to the endothelial secretome are critical for melanoma L-EV-induced angiogenic potential. These findings collectively show that EV subtypes behave differently in context of tumor type and they describe a unique mechanism of bevacizumab resistance in melanoma, thus providing further insight into the mixed clinical success of current therapeutics.

## RESULTS

### Melanoma L-EVs promote endothelial angiogenic phenotypes in a mechanism distinct from sEVs

These investigations were initiated by assessing the effects of the EV subtypes— L-EVs and sEVs, alongside the potent angiogenic factor VEGF, on endothelial tube formation. EVs were isolated as previously described and fractionated into L-EV or sEV populations using differential ultracentrifugation and high resolution iodixanol (Optiprep) density gradients (Figure S1A-D) (21). Consistent with previous reports, L-EV fractions mainly consisted of MVs based on size and protein composition including β-actin, ARF6, and Annexin A1, while the sEV fractions were smaller in size and enriched in exosome markers, ALIX and CD81 (21).

Endothelial tube formation was examined using a well-established in vitro assay for angiogenesis wherein endothelial cells organize to form capillary tube structures (22, 23). We used HUVECs (human umbilical vein endothelial cells) and SVEC4-10 mouse endothelial cells for these studies. To assess the effects of L-EVs, isolated vesicles were incubated with endothelial cells and resulting capillary networks were analyzed for tube features including the number of segments, nodes, junctions, meshes (24). We assessed whether L-EVs were taken up by recipient cells by monitoring in parallel, the uptake of fluorescent L-EVs isolated from the melanoma cell line, LOX, expressing GFP-tagged Annexin A1, an MV marker (21, 25). Fluorescent GFP puncta were visible in the recipient cell cytoplasm, indicating that L-EVs were internalized by endothelial cells (Figure S1E). These studies revealed that L-EV uptake promoted robust tube formation in a dose-dependent manner (Figure S1F) and enhanced elongation of tube networks relative to mock-treated cells, and comparable to that observed upon stimulation with VEGF alone (Figure 1A, 1B, S2A). Notably however, the melanoma sEV fraction had little to no effect on endothelial tube formation. To confirm that this finding was not exclusively a function of the cell line used, we examined the effect on EVs on the paired primary and metastatic melanoma cell lines, A375P and A375-MA2. We observed that L-EVs but not sEVs derived from both A375P and A375-MA2 melanoma cell lines increased tube formation (Figure 1C, 1D, S2B). Thus, the effects of EVs on endothelial tube formation appeared to be consistent across melanoma cell lines and independent of tumor stage.

**Figure 1.**
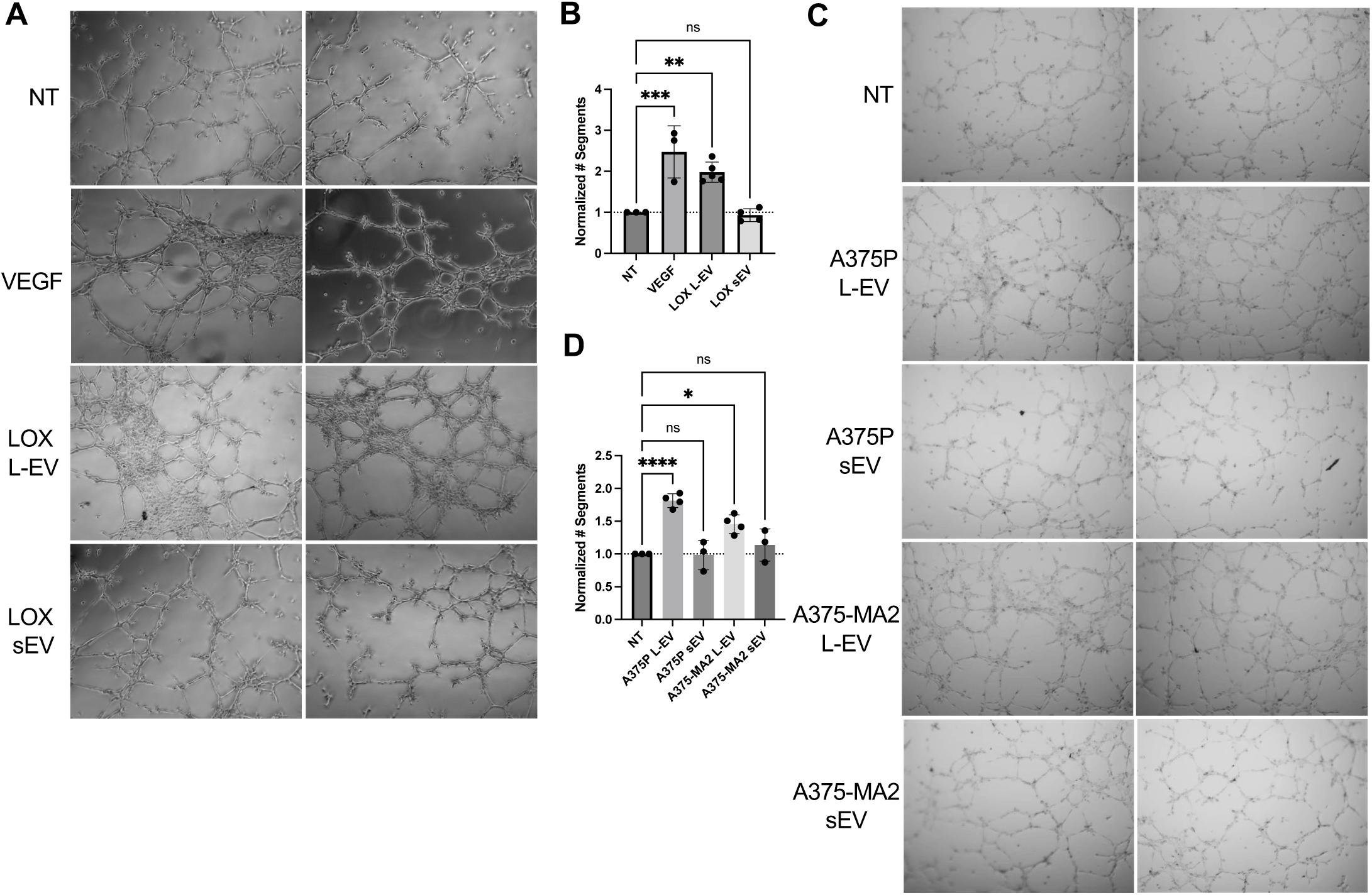
Melanoma L-EVs increase endothelial tube formation distinct from sEVs. (A) SVEC4-10 endothelial cells were incubated with no treatment 1XPBS control, VEGF (20ng/mL) or LOX L-EVs or sEVs, as described in the methods. Cells were allowed to form tubes and networks were imaged on an inverted microscope after 5 hours. (B) Resulting tube networks were analyzed for number of segments using Fiji and normalized to respective no treatment controls among replicates. (C,D) Endothelial cells were incubated with L-EVs or sEVs derived from paired primary and metastatic cell lines A375P or A375-MA2 and number of segments were quantified using Fiji. Data are presented as means ± SD. The p-values were obtained by one-way ANOVA with Dunnett’s correction (ns: not significant, *p<0.05, ** p<0.01, ***p<0.001, ****p<0.0001) from at least 3 independent experiments.

Given prior reports on the effects of sEVs on pro-angiogenic phenotypes in other tumor cell types (20, 26), we assessed the effects of large and small EVs on two additional tumor cell lines: MDA-MB-468, a metastatic breast cancer cell line; and 786-O, a clear cell renal carcinoma cell line. MDA-MB-468-derived L-EVs resulted in increased tube formation in similar ways to that described above for melanoma L-EVs (Figure S3A-C), whereas sEVs from the same cell line had no effect on tube formation. The effects of breast cancer cell-derived L-EVs in promoting pro-angiogenic phenotypes are consistent with prior reports (27). With respect to 786-O cells, however, both the L-EV and sEV fractions increased endothelial tube formation. Further, when compared to the effects of breast and melanoma cell types, the induction of nodes, junctions, and meshes were even stronger with 786-O L-EVs (Figure S3A-C). The sEV fraction from 786-O cells formed dense networks and a distinct sprouting morphology evident through increased branching. Endothelial branching or sprouting phenotypes are indicative of tip cell formation and are characterized by extensions at the edges of vascular sprouts (28). As such, both the 786-O small and large EV fractions enhanced pro-angiogenic phenotypes, although in unique ways.

These concerted and potent effects of renal EVs may contribute to the highly vascularized nature of ccRCC tumors (29). Notably, treatment of endothelial cells with an equivalent amount of EVs from a non-tumorigenic fibroblast cell line had no effect on tube formation, indicating that the angiogenic growth-inducing effects are specific to vesicles shed from the tumor cell lines (Figure S3D).

In light of the above findings, we examined melanoma L-EVs for the presence of pro-angiogenic cargos identified in EVs shed from various tumor cell types. We first examined for the presence of CD133, which has been shown to be a pro-angiogenic cargo of L-EVs-derived from colorectal cancer cells (30). CD133 was not present in the melanoma cells by western blotting of cell lysates (Figure S4A-B). We also examined for the presence of EphB2, which was reported to be pro-angiogenic cargo on sEVs derived from head and neck squamous cell carcinoma (HNSCC) cells (26). We observed that EphB2 was not expressed in the melanoma and MDA-MB-468 cell lines nor was it detectable in L-EV and sEV fractions (Figure S4C-D). However, it was expressed in the 786-O ccRCC cell line. Further, EphB2 appeared to be enriched in the 786-O sEV fraction (Figure S4E). Taken together, these studies show that the effects of EV subtypes on endothelial cells vary with tumor origin and that key pro-angiogenic cargos vary between EV subtypes. They also highlight the importance of understanding the distinct roles of EV subtypes in angiogenesis.

### Melanoma L-EVs contain VEGF as a luminal cargo

Given that VEGF is a major factor that drives angiogenesis, we investigated melanoma L-EVs for VEGF as a cargo using a combination of morphological and biochemical approaches. VEGF is thought to be present on EVs either as luminal or surface membrane-bound cargo (19, 20, 27). At the surface of EVs, VEGF is able to directly bind receptors on recipient cells, whereas luminal VEGF is shielded from direct interactions with recipient cells. We assessed vesicle morphology and localization of VEGF via confocal and STORM imaging. When L-EVs were processed for confocal imaging in the presence or absence of a membrane permeabilizing agent, we observed at 1.5 fold increase in labeling for VEGF-A in the presence of permeabilizing agent (Figure S5A). L-EVs were marked by labeling with β1 integrin, a transmembrane cargo, that remained constant under both experimental conditions. For super-resolution STORM imaging, melanoma L-EVs were marked with Pan-EV, a membrane dye, with and without permeabilization. VEGF labeling was barely detectable relative to that observed in the presence of permeabilizing agent (Figure 2A). These studies indicate that the majority of melanoma L-EV associated VEGF is present in the lumen. Similarly, when L-EV-associated VEGF was examined by ELISA using both intact and lysed L-EVs, the VEGF signal was higher in lysed L-EVs, indicating that the majority of VEGF was present in the L-EV lumen (Figure 2B). Finally, trypsin digestion to degrade surface-bound protein cargo had no effect on L-EV VEGF. As controls, Fascin, a known luminal cargo, was not sensitive to trypsin degradation under the same experimental conditions, whereas transmembrane cargos MMP14 and CD81 were sensitive to trypsin. In all cases, L-EV cargos were sensitive to enzymatic digestion in the presence of detergent (Figure 2C-D). Collectively, these results suggest that the majority of VEGF is in the lumen of melanoma L-EVs with only a relatively small pool of VEGF at the L-EV surface.

**Figure 2.**
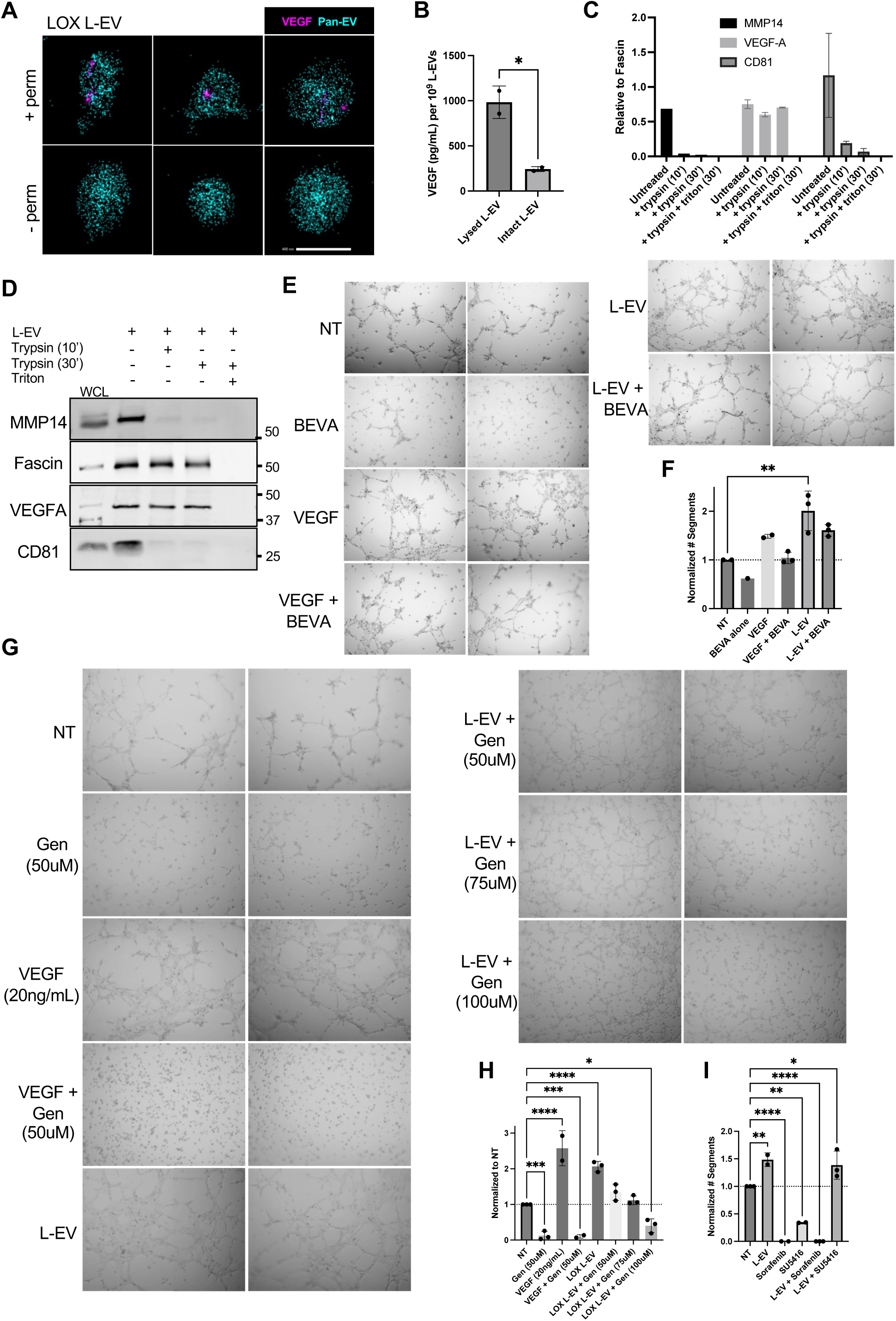
VEGF-A is primarily a luminal cargo of melanoma L-EVs. (A) L-EVs were purified from LOX melanoma cells as described in the methods and processed for super-resolution microscopy. EVs were captured using phosphatidylserine and stained with VEGF-A and Pan-EV in the presence (+ perm) or absence (-perm) of saponin. Scale bar, 400nm. (B) L-EVs were isolated and resuspended in 0.1µm filtered 1XPBS or in RIPA lysis buffer. Samples were used to perform a VEGF-A ELISA and concentrations per 10^9^ L-EVs were calculated based on a 4-parameter logistic (4PL) curve fit model generated from standard concentrations. (C,D) The same number of isolated L-EVs (10^9^) were resuspended in 0.1µm filtered 1XPBS with no treatment, 1X Trypsin-EDTA (10 minutes or 30 minutes, as indicated) or a combination of 1X Trypsin-EDTA and 0.1% Triton X-100. L-EVs were re-isolated, re-suspended in 1.5X loading dye, and separated by SDS-PAGE for western blot analysis. Pixel densities were quantified using Fiji and independent replicates were plotted as means ± SD. Molecular weight markers (kD) are indicated. (E,F) SVEC4-10 endothelial cells were incubated with no treatment vehicle control (NT), 100nM bevacizumab (BEVA), 20ng/mL VEGF, LOX L-EVs or combination treatments, as described in the methods. Tube networks were imaged after 5 hours and analyzed in Fiji for number of segments. Values among replicates were normalized to respective no treatment controls. (G) SVEC4-10 cells were incubated with 50-100μM genistein (Gen), 20ng/mL VEGF, LOX L-EVs, or combination treatments, as indicated. Tube networks were allowed to form and imaged after 5 hours. (H) Tube analysis was conducted in Fiji to determine the number of segments after normalization to the no treatment control condition. (I) SVEC4-10 cells were incubated with 10μM SU5416 or 20μM Sorafenib alone or in combination with L-EVs. Data are presented as means ± SD. The p-values were obtained by one-way ANOVA with (F) Tukey’s correction or (H,I) Dunnet’s correction or (B) an unpaired two-tailed t test (ns: not significant, *p<0.05, ** p<0.01, ***p<0.001, ****p<0.0001) from at least 3 independent experiments.

### Effect of VEGF neutralization on L-EV-induced phenotypes

Given the above findings, we investigated the impact of L-EVs in promoting endothelial angiogenic phenotypes in the presence of VEGF neutralizing antibodies. VEGF neutralizing antibodies, including bevacizumab, have been used in the clinic as anti-angiogenic therapies, but with mixed success in melanoma (31). In tube assays, addition of bevacizumab markedly reduced VEGF-induced tube formation to baseline levels (Figure 2E-F). Notably, under the same experimental conditions, melanoma L-EV-treated cells exhibited endothelial tube structures above baseline (Figure 2E-F). This decreased sensitivity to bevacizumab is somewhat analogous to a previous report in breast cancer cell lines which demonstrated that crosslinked VEGF on MVs interacts with HSP90 in a manner that renders breast cancer cell lines insensitive to bevacizumab (27). The study also showed that pretreatment with 17-AAG, an HSP90 inhibitor released VEGF and re-sensitizes the breast MVs to bevacizumab. However, we found that pre-treating melanoma L-EVs with 17-AAG had no effect on tube formation (Figure S5B-C).

Additionally, VEGF interactions with heparin-sulfate on the EV surface have also been implicated in bevacizumab-insensitivity (20). We ruled out this mechanism as heparinase treatment had no effect on melanoma L-EV-induced endothelial tube formation (Figure S5D-E). These studies show that VEGF is predominantly a luminal cargo of melanoma L-EVs, which potentially establishes a unique mechanism for bevacizumab insensitivity. Taken together, our findings suggest that there are multiple mechanisms by which EV-associated VEGF may confer bevacizumab-insensitivity.

To further investigate the effect of blocking VEGF signaling, endothelial cells were treated with a tyrosine kinase inhibitor, genistein. Genistein is known to inhibit endothelial tube formation (32), and consistently, treatment with 50uM genistein inhibited tube formation (Figure 2G-H). VEGF-induced tube formation was also blocked under the same experimental conditions.

However, the addition of genistein only partially arrested L-EV-induced tube formation. A decrease in tube thickness was observed alongside a dose-dependent decrease in the number of segments, but without significant effect on nodes, junctions, tube length or meshes (Figure 2G, 2H, S5F). At significantly higher concentrations of genistein (100uM), there was a marked loss of capillary networks although remnants of tube formation were still observed. These data suggest that L-EVs elicit additional signaling events to drive endothelial tube formation relative to VEGF alone that may involve tyrosine kinase-independent alterations to endothelial tube formation. To investigate this contention further, we examined the effects of SU5416, a selective inhibitor of VEGFR2, and Sorafenib, a multi-kinase inhibitor with potent inhibitory effects on Raf kinases as well as VEGFR2/3, PDGFRβ, FLT3, and c-Kit (33). Treatment with Sorafenib alone blocked tube formation, whereas treatment with SU5416 alone resulted in fewer tubes and a partial block in segment formation (Figure 2I, S5G). However, the addition of SU5416 had no effect on L-EV stimulated tube formation, supporting the above findings that the direct effects of melanoma L-EVs are independent of VEGFR. In contrast, treatment with Sorafenib inhibited L-EV-induced tube formation, suggesting that melanoma L-EVs may act through activation of Raf kinases or one of the aforementioned receptors.

### Melanoma L-EVs facilitate autocrine endothelial VEGF signaling

To gain insight into the mechanisms by which melanoma L-EVs promote endothelial capillary tube formation, we first examined the effect of L-EVs on endothelial VEGF production. Endothelial cells were incubated with melanoma L-EVs for 48 hours, EVs were sedimented by centrifugation, following which conditioned media was examined for soluble VEGF by ELISA. We observed a 5.4-fold increase in soluble VEGF after treatment with melanoma L-EVs (Figure 3A). VEGF release was dose-dependent (Figure 3B) and evident as early as 3 hours post L-EV treatment (Figure 3C). To determine whether the increased VEGF release involved transcriptional regulation, we examined VEGF mRNA by qRT-PCR. VEGF mRNA increased only at the later time points i.e. 24 and 48 hours, post L-EV treatment (Figure 3D).

**Figure 3.**
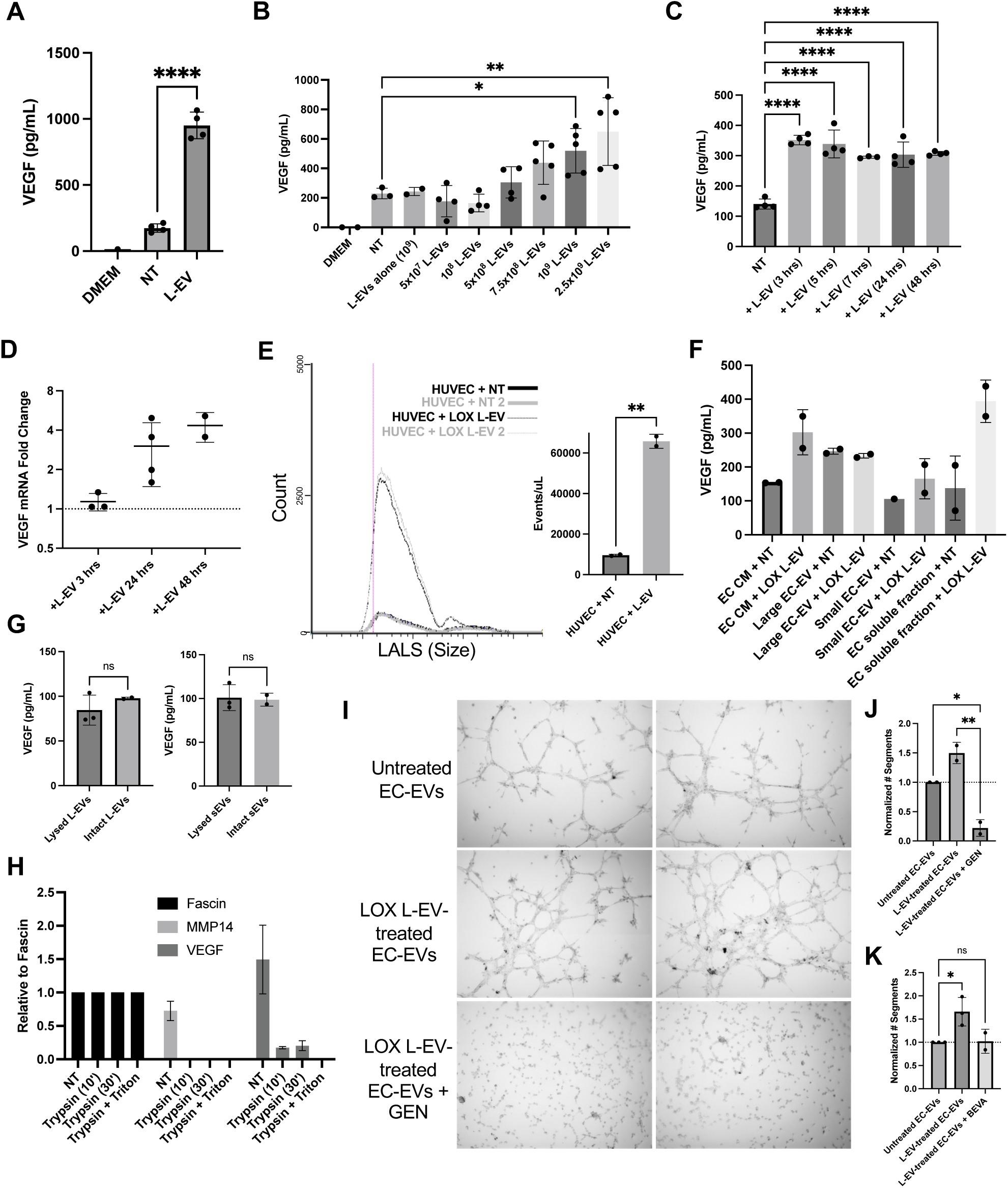
Melanoma L-EVs alter endothelial VEGF release. (A) HUVEC cells were treated with 1XPBS control (NT) or LOX L-EVs for 48 hours. Conditioned media was collected and EVs were removed by ultracentrifugation. Remaining supernatant was used for VEGF ELISA analysis. Concentrations were calculated based on a 4PL curve fit model generated from standard concentrations. (B) HUVEC cells were treated with varying amounts of LOX L-EVs, as indicated. Supernatants were depleted of EVs and used for VEGF ELISA analysis. (C) HUVEC cells were treated with equivalent amounts of LOX L-EVs for various timepoints, as indicated. Supernatants were depleted of EVs and used for VEGF ELISA analysis. (D) Total RNA was isolated from HUVECs after treatment with equivalent amounts of LOX L-EVs for indicated times and used to perform qRT-PCR on VEGF-A, as described in the methods. Experiments were repeated at least 3 times and outliers were removed. (E) HUVEC cells were treated with LOX L-EVs or 1XPBS control (NT) for 16 hours. Media was removed and ECs were allowed to shed into EV-depleted media for 72 hours. Total EC-EVs were collected and analyzed on a microparticle flow cytometer. Particle count vs large angle light scatter (LALS) was plotted and particle events per microliter were used to generate a bar graph. (F) HUVEC cells were treated with LOX L-EVs or 1XPBS (NT) for 16 hours and media was replaced with EV-free. Conditioned media (CM) was fractionated and subject to VEGF ELISA analysis. VEGF concentrations were calculated based on a 4PL curve fit model generated from standard concentrations. (G) L-EVs and sEVs were purified from LOX L-EV-treated or untreated endothelial cells by serial ultracentrifugation and resulting EV populations were either lysed in RIPA buffer (“lysed”) or maintained in PBS (“intact”) and subject to a VEGF ELISA. (H) EC-EVs were collected by ultracentrifugation and used to perform a trypsin digestion as outlined in the methods. EC-EVs were re-isolated, re-suspended in 1.5X loading dye, and separated by SDS-PAGE for western blot analysis. Pixel densities were quantified using Fiji and normalized relative to Fascin. (I-K) EC-EVs were collected by ultracentrifugation as outlined above from endothelial cells treated with 1XPBS (untreated) or LOX L-EVs. EC-EVs were used to perform a tube assay in the presence (or absence) of genistein or bevacizumab, as indicated. Tube networks were analyzed after 5 hours in Fiji. Independent biological replicates are plotted as means ± SD. The p-values were obtained by one-way ANOVA with (J) Tukey’s correction or (A-C,K) Dunnet’s correction or (E,G) an unpaired two-tailed t test (ns: not significant, *p<0.05, ** p<0.01, ***p<0.001, ****p<0.0001).

To determine if melanoma L-EV treatment impacts endothelial cell-EV (EC-EV) release, HUVECs were incubated with melanoma L-EVs followed by microflow cytometry to examine vesicle release. These studies revealed an increase in overall EC-EV shedding (Figure 3E). ELISA analysis of the fractionated conditioned media for VEGF showed that while a significant portion of the L-EV-mediated increase in VEGF secretion is soluble VEGF (Figure 3F), endothelial EVs, both L-EVs and sEVs, also contain VEGF as a surface cargo (Figure 3G). Super-resolution imaging of EC-EVs labeled for VEGF-A with or without permeabilization, showed that VEGF is present at the surface of EC-EVs, both large and small (Figure S6A). Likewise, trypsin digestion experiments on the EC-EVs showed that, similar to MMP14, surface VEGF was sensitive to trypsin treatment (Figure 3H, S6B).

Next, we assessed tube formation with EVs derived from untreated or LOX L-EV-treated endothelial cells. To this end, EVs released from the same number of endothelial cells were collected. Likely due to increased numbers of EVs, EC-EVs collected from LOX L-EV-treated cells significantly increased all tube features including number of segments, meshes, nodes, and junctions relative to EVs released from untreated endothelial cells (Figure 3I, 3J, S6C). Notably, EC-EVs collected from untreated endothelial cells also enhanced segment formation, though to a lesser extent than L-EV-treated EC-EVs (Figure S6D). In addition, genistein treatment of EC-EVs markedly inhibited the effects of EC-EVs (Figure 3I-J). We also tested the effect of bevacizumab with EC-EVs and found that VEGF on the surface of EC-EVs was sensitive to bevacizumab (Figure 3K, S6E). Thus, these results suggest that melanoma L-EVs trigger endothelial release of soluble VEGF, as well as EC-EVs with VEGF at the surface, thereby enabling tyrosine kinase-dependent autocrine signaling to drive endothelial tube formation.

Taken together, the studies described thus far suggest that melanoma L-EVs contain VEGF as a luminal cargo and induce sustained endothelial tube formation through mechanisms that are both, tyrosine kinase dependent and independent. Tyrosine kinase dependent-mechanisms involve altering endothelial cell secretion of both soluble VEGF and EC-EV surface-associated VEGF. Additionally, combination treatment of genistein with melanoma L-EVs suggest that mechanisms independent of tyrosine kinase regulation contribute to pro-angiogenic phenotypes.

### Melanoma L-EVs alter the angiogenic cytokine profile of endothelial cells

We investigated other VEGF-independent modes by which L-EVs promote endothelial tube formation. Prior literature demonstrated a role for cytokines in promoting angiogenesis via mechanisms that are both VEGF-dependent and independent (34). Interleukins, including IL-1, IL-8 and IL-6, are able to induce expression of VEGF as well as directly induce angiogenic phenotypes such as proliferation, tube formation, survival and matrix metalloproteinase production (35–37). We profiled cytokines released by L-EV-treated endothelial cells. To this end, endothelial cells were treated with melanoma L-EVs or mock treated for 48 hours, cellular RNA was collected followed by cytokine profiling quantitative PCR. Independent biological replicates revealed increased endothelial expression of interleukins IL-12α, IL-18, IL-1 (α/β), IL-6 and IL-8 (Figure 4A) and decreased expression of IFNα6 and IL-16. The alterations in cytokine transcripts were validated by analyzing cytokine and chemokine release from L-EV-treated endothelial cells using a human cytokine antibody array (Figure 4B-C). Independent biological replicates showed increased levels in the culture media of IL-8, macrophage migration inhibitor factor (MIF), and plasminogen activator inhibitor (Serpin E1/PA-I), all of which are pro-angiogenic. Further, to assess that effects of released cytokines on endothelial tube formation, we performed a tube assay with L-EV-depleted endothelial cell conditioned media in the presence (or absence) of a MIF neutralizing antibody. Neutralizing MIF resulted in a reduction in all tube network features (Figure 4D-E, S7) relative to control experimental conditions. This proof-of-principle experiment demonstrates that melanoma L-EVs drive endothelial tube formation in part by regulating endothelial cytokine release.

**Figure 4.**
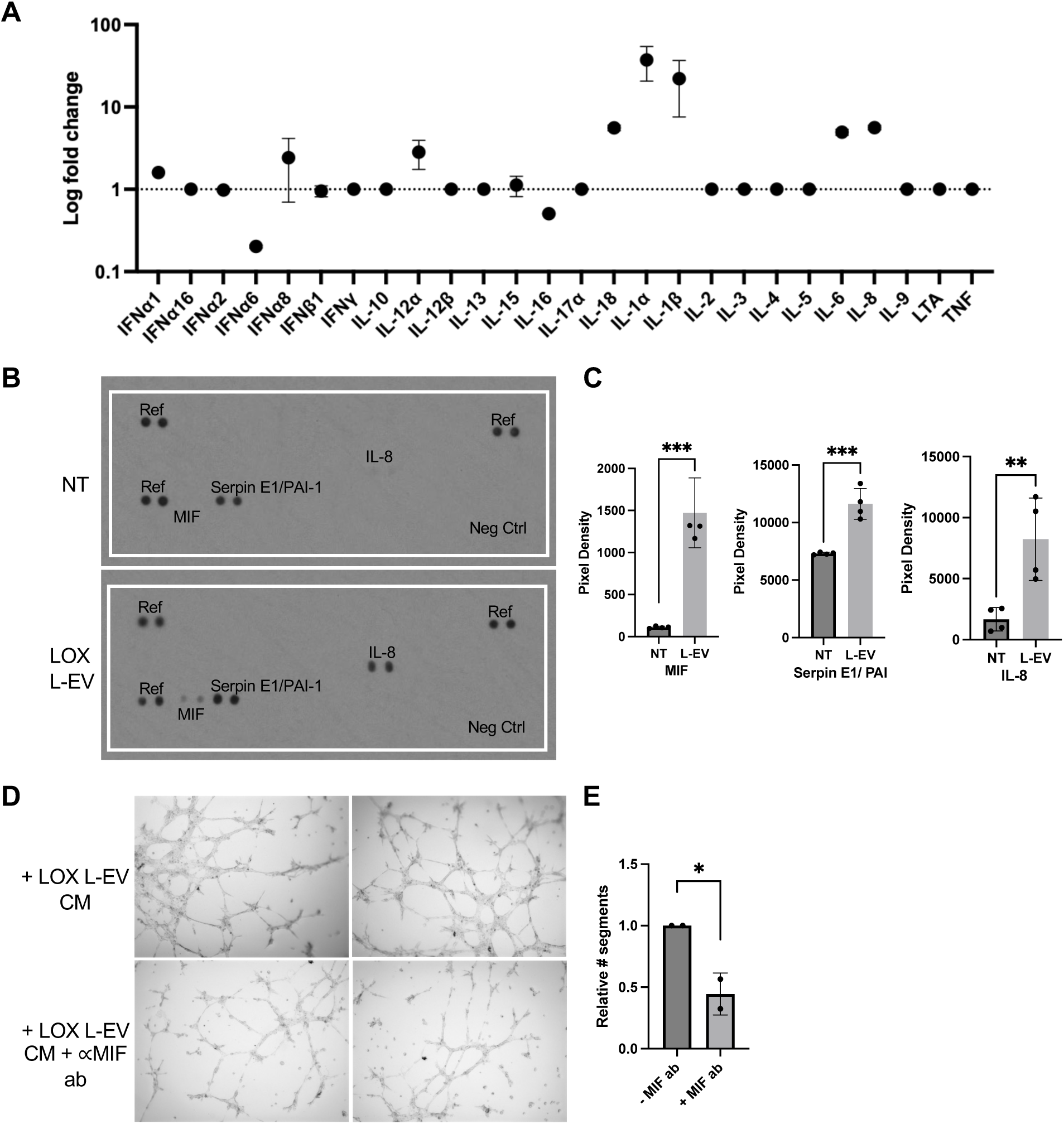
L-EV-treated ECs exhibit a distinct cytokine profile. (A) HUVEC cells were treated with LOX L-EVs or 1XPBS control for 48 hours and RNA was collected to perform a qRT-PCR cytokine array. Comparative C_T_ analysis was performed using an 18S endogenous control and relative expression of each cytokine was plotted. The data are presented as means of two independent replicates. (B) HUVEC cells were treated as described with either 1XPBS or LOX L-EVs and conditioned media was collected. EVs were removed and remaining culture media was used to perform a cytokine dot blot array. (C) Pixel density analysis of the dot blot array was performed using Fiji, and values were normalized to reference dots. Independent replicates are plotted. (D,E) Endothelial cells were treated with LOX L-EVs for 16 hours and EVs were removed. Media was replaced with EV-free and cells were incubated for 48 hours. Conditioned media was used in combination with a MIF neutralizing antibody (ab) (5µg/mL) to perform a tube assay. Data are presented as means ± SD. The p-values were obtained by an unpaired two-tailed t test (ns: not significant, *p<0.05, ** p<0.01, ***p<0.001, ****p<0.0001).

## DISCUSSION

The distinctive mechanisms of cell-to-cell communication offered by small and large EVs are attracting significant interest in various biological contexts and disease states including cancer. EVs contribute to cancer progression in various ways and are being explored as targets for new cancer therapies (38, 39). Both L-EVs and sEVs have been suggested to exert major roles in this regard. For instance, EVs have been thought to exert significant influence on tumor angiogenesis, with previous reports showing that one or more non-overlapping cargos contained in the shed vesicles from various cell types promote pro-angiogenic phenotypes (40, 41). Here, we report on a unique mechanism by which L-EVs, but not sEVs, shed from melanoma cells induce a robust and sustained response in endothelial cells triggering endothelial tube formation that is insensitive to the inhibitory actions of bevacizumab, an anti-VEGF therapeutic as well as SU5416, a VEGF receptor inhibitor. However, the induction of angiogenic phenotypes were sensitive to sorafenib, a multi-kinase inhibitor. L-EV-induced paracrine stimulation modulates sustained endothelial autocrine behaviors by augmenting the secretion of pro-angiogenic cytokines and endothelial cell-derived extracellular vesicles (Figure 5). Finally, we show that the effects of tumor EV subtypes on endothelial cells vary with tumor origin and that key pro-angiogenic cargos can vary between EV subtypes and across tumor cell lines. These findings are noteworthy in that they elucidate a mechanism by which L-EVs facilitate the induction of pro-angiogenic phenotypes and provide potential insight into the limited efficacy of bevacizumab in melanoma. Moreover, they also emphasize the importance of understanding the distinct roles of EV subtypes in angiogenesis and highlight the potential of simultaneously targeting the endothelial secretome as a complement to therapeutic strategies for advanced melanoma.

**Figure 5.**
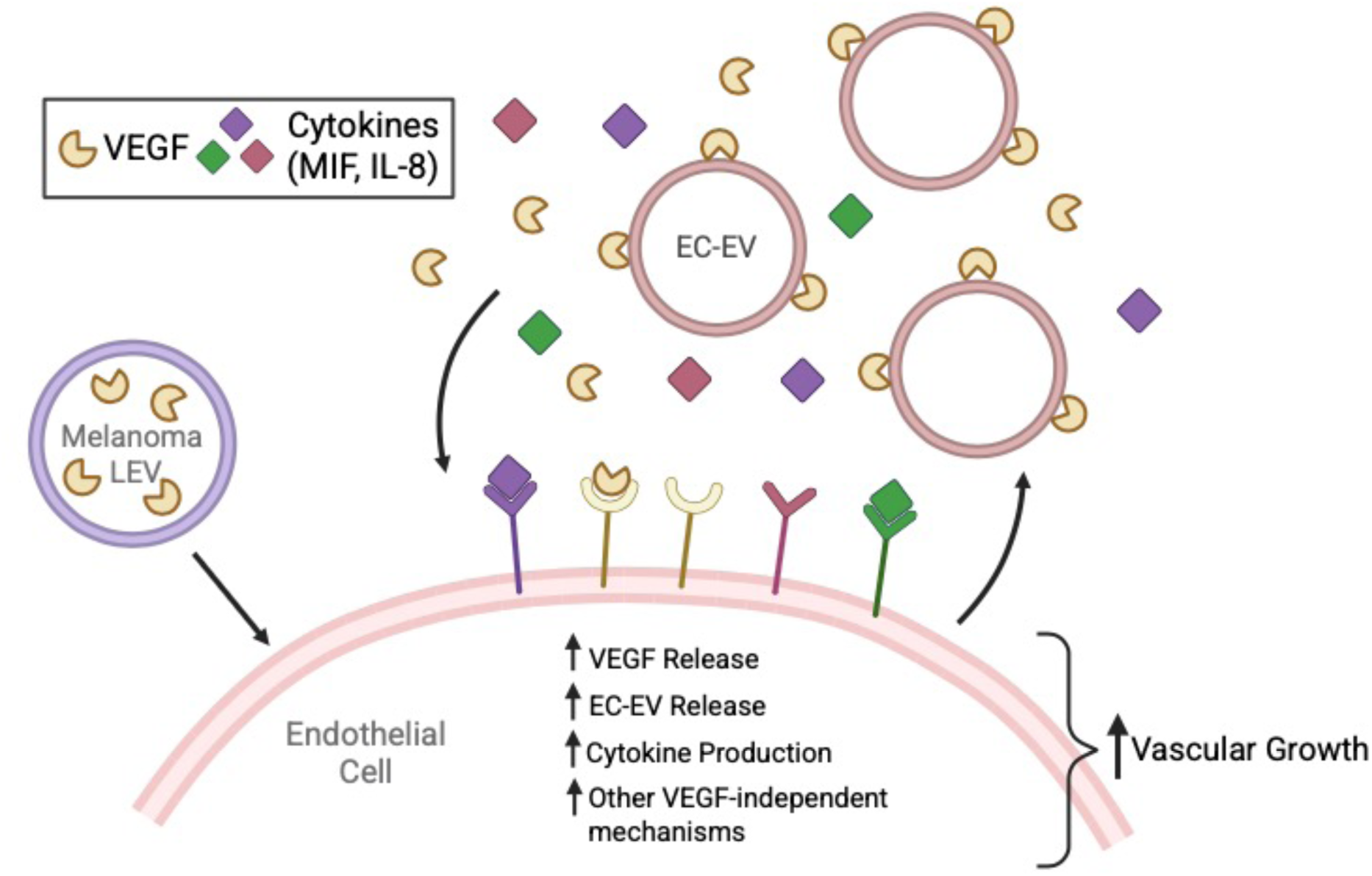
Schematic depicting the paracrine and autocrine effects of melanoma L-EVs on endothelial angiogenic phenotypes. Melanoma L-EVs, containing VEGF as a luminal cargo, facilitate paracrine signaling by inducing the secretion of soluble VEGF, EC-EVs, and cytokines from endothelial cells, all of which participate in sustained autocrine signaling to drive vascular growth.

The therapeutic targeting of VEGF to inhibit tumor angiogenesis has yielded mixed outcomes in melanoma. For instance, a retrospective study conducted on advanced melanoma patients indicated that employing bevacizumab as a first- or second-line therapy resulted in limited success, with objective response rates of 6.3% and 3.4%, respectively (42). Other studies have demonstrated limited efficacy when bevacizumab is used alone, but enhanced outcomes when combined with interferon or chemotherapy (31, 43). These clinical studies on bevacizumab’s efficacy in melanoma underscore the importance better understanding the molecular basis of bevacizumab-resistance.

Large EVs released by melanoma lines tested here are predominantly MVs based on particle size of the isolated L-EV fraction, however, we do not exclude the potential contribution of other even larger L-EV subtypes (44) that may be also present in low abundance. Of note, these pro-angiogenic EC-EVs by most measures are distinct from the organelle-rich apoptotic body or exopher-like large endothelial-derived EVs greater than 1 micron that have been shown to promote vasculature damage (45). We demonstrate that melanoma L-EVs harbor VEGF as a luminal cargo, which is shielded from anti-VEGF therapeutic antibodies. Furthermore, inhibition of tyrosine kinase activity resulted in a diminution in endothelial tube formation, indicating that L-EVs induce kinase-dependent, yet VEGF-independent endothelial activation. L-EV induced tube formation was also sensitive to Sorafenib treatment. Sorafenib, a multi-kinase inhibitor has been shown to target endothelial cells by disrupting various pro-angiogenic pathways including the receptor tyrosine kinases VEGFR-2/3 and PDGFR-β as well as the Raf-MEK-ERK and PI3K-AKT pathways with inhibitory effects on cell proliferation, migration, and new vessel formation (33). The drug also affects other pathways like JNK with impact on cell survival. In future work it will be of interest to determine which of these pathways are sensitive to Sorafenib downstream of L-EVs.

In addition to the studies we describe here, previous reports have also indicated that EVs protect vascular endothelial growth factor (VEGF) from therapeutic detection. For example, VEGF isoforms present on the surface of EVs derived from breast tumor and oral squamous carcinoma cell lines among others, are resistant to bevacizumab treatment, where depending on the tumor type, interactions between VEGF and EVs mediated by heparan sulfate or HSP90 were reported to be responsible for bevacizumab insensitivity (19, 20, 27). However, here we found that blocking the effect of heparan sulfate or HSP90 had no impact on melanoma L-EV-induced tube formation. We also note that melanoma L-EV-stimulated endothelial cells released soluble VEGF as well as EVs bearing VEGF on their surface, which were responsive to tyrosine kinase inhibition and bevacizumab treatment. Despite this, the paracrine effects of melanoma L-EVs are insensitive to bevacizumab treatment, pointing to the complexity of bevacizumab sensitivity. Thus, while the luminal positioning of VEGF likely significantly contributes to bevacizumab insensitivity other mechanisms including enhanced cytokine release, as discussed below, may also be responsible.

In addition to VEGF secretion, our studies also indicate that L-EVs markedly contribute to the augmentation of soluble cytokine release namely, MIF, IL-8, and PAI-1 from endothelial cells. MIF is known to facilitate tumor progression and angiogenesis by inducing endothelial proliferation and in vivo blood vessel formation (46, 47). IL-8 is another well-established pro-angiogenic factor that directly influences the proliferation, migration, and tube formation of endothelial cells (35, 48). In fact, in addition to elevated levels of VEGF and other growth factors, IL-8 secretion is also associated with high tumor burden, advanced disease stage, and poor overall survival in melanoma (49). HuMax-IL8, which inhibits IL-8, has been shown to enhance clinical outcomes in gastric and NSCLC tumors by reducing serum IL-8 levels and sensitizing tumors to other immunotherapies, including PD-1/PD-L1, although there are no reports on IL-8 inhibition on clinical outcomes in melanoma (50, 51). The uPAR/uPa system is also recognized as a regulator of angiogenesis, activating MMPs to degrade extracellular matrix and release matrix-bound factors such as vascular endothelial growth factor (VEGF) (52). Thus L-EVs may also indirectly promote angiogenesis by altering the uPAR system. We note that the expression of several cytokines including IL-12α, IL-18, IL-1 (α/β), and IL-6 were increased in response to melanoma L-EVs. It is possible that some of these cytokines may also be EV-associated although this remains to be ascertained. Notably in this regard, serum cytokine levels in melanoma are often correlated with poor prognosis and late disease stage (53). Increased IL-6 is a prognostic factor for shorter overall survival in advanced melanoma patients receiving immune checkpoint inhibitors or chemotherapy (54).

There is much that remains to be understood regarding the mechanisms by which melanoma L-EVs contribute to tumor angiogenesis. Areas of future research include identifying the surface molecules essential for L-EV-endothelial cell interactions and cargos responsible for promoting angiogenic phenotypes. Additionally, the activation of VEGF-dependent and independent signaling pathways in endothelial cells in response to L-EVs warrants further investigation.

Finally, investigating the efficacy of sorafenib therapies or the synergistic effects of blocking cytokines with anti-angiogenic therapy in models of metastatic melanoma, where bevacizumab efficacy demonstrates variability, could provide valuable insights. Although the answers to these and other questions surrounding L-EV function remain elusive, it is evident that the actions of L-EVs and other EVs in tumor progression hold significant promise as exciting research areas with the potential to profoundly impact future translational efforts.

## DATA AVAILABILITY

The data supporting the findings of this study are available within the article and its supplemental information. Requests for additional information may be directed to the corresponding author.

## SUPPORTING INFORMATION

This article contains supporting information.

## ACKNOWLEDGEMENTS

We thank Regan Moore from ONI, and Alex Boomgarden, Madison Schmidtmann, Arushi Bute, and Marie Gerges in the D’Souza-Schorey lab, for their helpful contributions to this study. This work was supported in part by NIH 1R01GM148708, NIH 1R03CA273598-01 and the Boler Foundation to CD-S. The content is solely the responsibility of the authors and does not necessarily represent the official views of the National Institutes of Health.

## CONFLICT OF INTEREST DISCLOSURE

The authors declare that they have no conflicts of interest with the contents of this article.

## EXPERIMENTAL PROCEDURES

### Cell lines

LOX melanoma cells were previously described (Fodstad, et al., 1988) and kindly provided by Prof. Oystein Fodstad, Oslo University, Norway. A375P (RRID: CVCL_6233), A375-MA2 (RRID: CVCL_X495), MDA-MB-468 (RRID: CVCL_0419), 786-O (RRID: CVCL_1051), SVEC4-10 (RRID: CVCL_4393), HUVEC (RRID: CVCL_9Q53) and BJ Fibroblast (RRID: CVCL_3653) cell lines were purchased from ATCC. LOX were cultured in RPMI supplemented with 10% Atlas serum, 2 mM L-glutamine and 100 U/mL penicillin-streptomycin, 786-O were cultured the same as LOX cells with the addition of 1mM sodium pyruvate. A375P, A375-MA2, MDA-MB-468, SVEC4-10, BJ fibroblast cells were cultured in DMEM supplemented with 10% Atlas serum, 2 mM L-glutamine, 1mM sodium pyruvate and 100 U/mL penicillin-streptomycin, SVEC4-10 with heat-inactivated serum. HUVECs were cultured per manufacturer’s instructions in endothelial growth media (Lonza cat #CC-3162). All cell lines were routinely checked to ensure absence of mycoplasma contamination and maintained at 37°C in a humidified chamber containing 5% CO_2._

### Antibodies and other reagents

Antibodies to β1 integrin (AIIB2) and CD133 (0.5ug/mL) were purchased from the Developmental Studies Hybridoma Bank at the University of Iowa; antibodies to ALIX, β actin, VEGF-A, CD81 and ARF6, were purchased from Protein Tech; antibodies to β1 integrin (1952), MMP14, and Fascin were purchased from Millipore. EphB2 antibody was purchased from Cell Signaling. Pan-EV stain used for STORM imaging was from ONI. Secondary Alexa Fluor Plus antibodies and Phalloidin for immunofluorescence and western blotting were purchased from ThermoFisher. All antibodies were purchased from commercial sources and have been extensively validated by the manufacturer. Western blot analysis confirmed antibody specificity showing bands at expected molecular weights. Relevant positive and negative control lysates were used.

### Extracellular Vesicle Isolation

Cells were expanded to reach ∼80% confluency when growth media was replaced with media supplemented with EV-depleted serum (Gibco, ThermoFisher) for up to 3 days and media was collected. EVs were isolated by serial ultracentrifugation and extensively washed with PBS and/or further purified using an iodixanol density gradient, as previously described (21, 25). All EV fractions were analyzed for particle concentration and size using a micro flow cytometer (Apogee Flow Systems). Calibration gates were determined using a mixture of silica beads at 180, 240, 300, 590, 880, and 1300 nm diameters. Data analysis was done using the Histogram Software (Apogee Flow Systems).

### Endothelial Tube Formation Assay

For the endothelial tube formation assay, SVEC4-10 endothelial cells (3×10^4^) were plated onto low growth factor Matrigel (Corning) in serum-free DMEM with 2% heat-inactivated FBS plus either extracellular vesicles (5×10^6^-1×10^9^), VEGF-A (20ng/mL), or genistein (50-100μM). Endothelial tube formation was monitored at 37°C and imaged on a Zeiss Axio Observer. Analysis of tube networks was conducted using the Angiogenesis Analyzer ImageJ plugin (24). For SU5416 and Sorafenib experiments, L-EVs were added to tube assays in combination with the indicated drugs at 10μM and 20μM concentrations, respectively. Where indicated, L-EVs were pre-treated with 100nM bevacizumab (MCE) or 0.9mU/mL Bacteroides Heparinase I (New England Biolabs) at 37°C. L-EVs were re-isolated and added to tube assays described above. For assays involving MIF neutralization, endothelial cells were incubated with isolated L-EVs, media was replaced and conditioned media was collected after 48 hours and used in combination with MIF neutralizing antibody (5μg/mL; R&D Systems) in tube assays. For 17-AAG treatments, LOX cells were treated with 17-AAG inhibitor (500nM-1μM; MCE) or an equivalent amount of DMSO in EV-free media and cells were allowed to shed for 24 hours before L-EV isolation. For combined 17-AAG and bevacizumab treatment, L-EVs were pre-treated with 17-AAG (10μM) for 90 minutes at 37°C and then used in combination with bevacizumab (0.5μg/mL) in tube assays. All treatment conditions were normalized to a no treatment vehicle control independently for each experiment.

### Western Blotting

Cells were lysed in RIPA buffer containing 25mM Tris-HCl pH 7.6, 150mM NaCl, 1% sodium deoxycholate, 0.1% SDS, 1% Triton X-100. Immediately prior to use, 1X phosphatase inhibitor and mammalian protease inhibitor cocktail (Millipore Sigma) were added. Lysates were processed for protein separation by SDS-PAGE and transferred to an immobilon-FL PVDF membrane (Millipore Sigma). Membrane blocking was performed in either 5% nonfat milk or 5% bovine serum albumin (BSA; Millipore Sigma) in TBS for 1 hour at room temperature.

Primary antibodies (diluted per manufacturer’s instructions) were incubated in 2% nonfat milk or BSA in TBS + 0.2% Tween-20 overnight at 4°C. Membranes were washed in TBS + 0.1% Tween-20, incubated with Alexa Fluor Plus secondary antibodies diluted 1:10,000 in 2% nonfat milk or BSA in TBS + 0.2% Tween-20 for 1 hour at room temperature in the dark. Membranes were again washed in TBS + 0.1% Tween-20 and then in TBS alone. A LiCor Odessey Scanner was used to image the blots, and pixel densitometry was performed in Fiji. COLO205 lysates were provided by cell signaling as a positive control.

### Trypsin digestions

EVs were incubated in 100µL 0.1µm 1X PBS with 1X Trypsin-EDTA or 0.1% Triton X-100 (final concentrations) for 10-30 min at 37°C. Enzymes and detergents were diluted by the addition of 900µL of 0.1µm 1X PBS. EVs were re-isolated by centrifugation.

### Immunofluorescence

Cells were plated on glass coverslips or EVs were plated on poly-L-lysine (MilliporeSigma) coated coverslips and allowed to adhere at 37°C before fixing in 2% paraformaldehyde (Electron Microscopy Supply). Coverslips were washed 3 times with PBS plus glycine and blocked/permeabilized in 5% BSA with 0.2% Triton-X-100 and 0.05% Tween 20. For experiments without permeabilization, blocking solution only contained 5% BSA. Coverslips were incubated in primary antibodies and further processed for immunofluorescence microscopy as previously described (21). To monitor L-EV uptake, cells were incubated with labeled L-EVs (isolated from the growth media of cells expressing GFP-tagged annexin A1). Coverslips were imaged on a Leica Stellaris 8 DIVE confocal microscope using the 60X and 100X objectives.

For super-resolution STORM microscopy, EVs were captured on prepared assay chips (EV Profiler Kit, Oxford Nanoimaging, UK) and stained with VEGF-A and Pan-EV in the presence or absence of permeabilizing agent (saponin). Samples were imaged with the ONI Nanoimager, Oxford Nanoimaging, UK. For EC-EVs, particle sizes were analyzed and EC-EVs sized <200 nm were binned as small EVs and >200 nm were binned as large EVs.

### Cytokine Array

Conditioned media from HUVEC cultures was harvested, spun at 100,000g, and the supernatants were tested for cytokine/chemokine levels using the Proteome Profiler Human Cytokine Array Kit (R&D Systems, Minneapolis, MN) according to the manufacturer’s instructions. Briefly, supernatants were mixed with a cocktail of biotinylated detection antibodies and incubated on nitrocellulose membranes that were pre-coated with 36 capture antibodies in duplicates.

Cytokine-antibody complexes were immobilized by the capture antibody on the membrane and unbound material was washed off. Membranes were incubated with streptavidin-HRP followed by chemiluminescent detection (ThermoFisher). Membranes were exposed to X-ray film and dot intensities were measured using Fiji.

### VEGF ELISA

For ELISA analysis, conditioned media from HUVEC cultures was tested for VEGF-A by ELISA (ThermoFisher) per the manufacturer’s instructions. Cell culture media was also tested alone to account for background levels of VEGF.

### Quantitative PCR

RNA was isolated from HUVEC cultures followed by reverse transcription (RT) using the TaqMan Fast Advanced Cells-to-C_T_ kit (ThermoFisher) per manufacturer’s instructions. Quantitative PCR (qPCR) was performed on a QuantStudio 5 real-time PCR machine (Applied Biosystems) and analyzed by the comparative C_T_ method. Amplification was measured using TaqMan gene expression assays for VEGF-A and 18S (ThermoFisher). Cycling conditions were set according to the manufacturer’s instructions. In separate experiments, cDNA was generated from HUVECs using the TaqMan Fast Advanced Cells-to-C_T_ kit (Invitrogen). qPCR was performed on a Step-One-Plus real-time PCR machine (Applied Biosystems) using a TaqMan Array Human Cytokine Network plate (Applied Biosystems) per manufacturer’s instructions. Briefly, cDNA was mixed with TaqMan Fast Advanced Master Mix and added to a plate that was pre-coated in 28 cytokine and 4 endogenous control assays. Reactions were cycled according to the manufacturer’s instructions and analyzed by the comparative C_T_ method.

### Statistical Analysis

Statistical analysis was performed using GraphPad Prism 10 version 10.4.2 or Microsoft Excel version 16.97.2. Student’s t-test was used to compare two groups with data that appeared to be normally distributed with similar variances. When comparing multiple treatment groups to a single control and the data were normally distributed, we performed one-way ANOVA with Dunnett’s correction for multiple comparisons. When multiple groups were analyzed and each group was compared to all other groups and the data appeared normally distributed, we used one-way ANOVA with Tukey test to correct for multiple comparisons. Experiments were performed with at least 3 biological replicates under similar conditions. For analysis of tube formation assay, treatment conditions were normalized to the no treatment vehicle control for each assay independently. For imaging experiments, representative images are shown with conclusions drawn from at least 10 fields of view across replicates. Additional statistical details are provided in figure legends.

## Abbreviations

**ccRCC:** Clear cell renal cell carcinoma; **EC-EV:** Endothelial cell-extracellular vesicle; **ELISA:** Enzyme-linked immunosorbent assay**; EV:** Extracellular vesicle**; HNSCC:** Head and neck squamous cell carcinoma; **HUVEC:** Human umbilical vein endothelial cell;**IL-**Interleukin-**MIF:** Macrophage migration inhibitor factor; **MV:** Microvesicle; **PAI-1:** Plasminogen activator inhibitor-1; **TME:** Tumor microenvironment; **VEGF**: Vascular endothelial growth factor

## Supporting Information

**Supplemental Figure 1.**
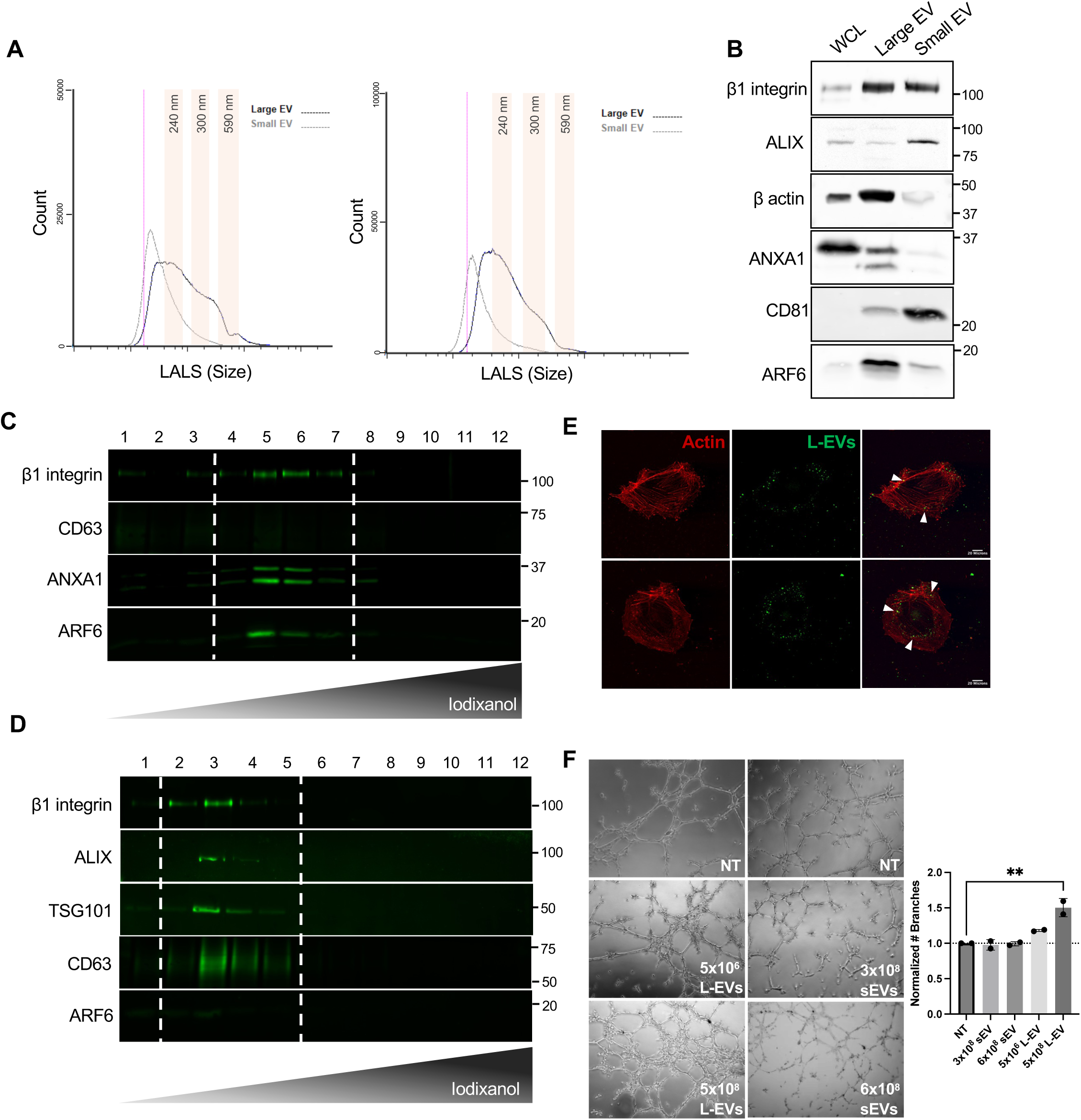
(A) L-EVs and sEVs were isolated from the conditioned media of LOX melanoma cells by serial ultracentrifugation and fractions were run on a micro particle flow cytometer. Histogram plots were generated as large angle light scatter (LALS) vs particle count and representative plots are shown. (B) Equal amounts of EV and whole cell lysates (WCL) were separated by SDS-PAGE and cargo contents were analyzed by western blotting. Molecular weight markers (kD) are indicated. (C,D) Isolated L-EV (C) or sEV (D) populations were purified on an iodixanol density gradient, as described in the methods, and contents were analyzed by western blotting for EV markers. (E) ANXA1-GFP tagged LOX L-EVs were incubated with HUVEC cells for 48 hours and cells were fixed and stained with phalloidin. Confocal microscopy reveals uptake of GFP-L-EVs by ECs with white arrows demonstrating internalized vesicles. (F) Isolated L-EV and sEV populations were used in a tube formation assay. The number of branches was quantified and normalized to the no treatment control. Values were plotted as means ± SD and the p-values were obtained by one-way ANOVA with Dunnett’s correction. (ns: not significant, *p<0.05, ** p<0.01, ***p<0.001, ****p<0.0001).

**Supplemental Figure 2.**
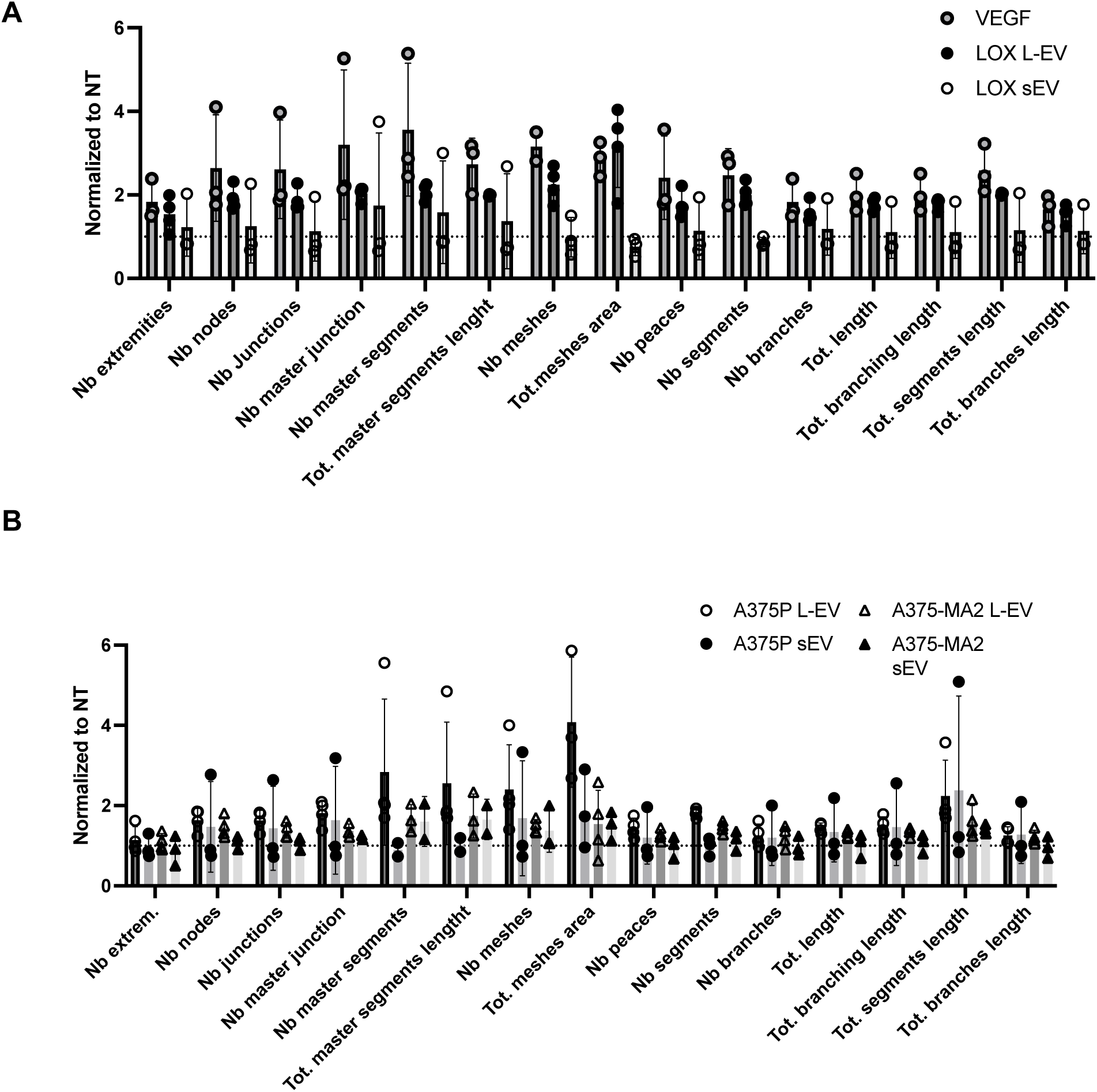
(A,B) Full network analyses for tube assays treated with melanoma L-EVs and sEVs. Data are presented as means ± SD. n≥3.

**Supplemental Figure 3.**
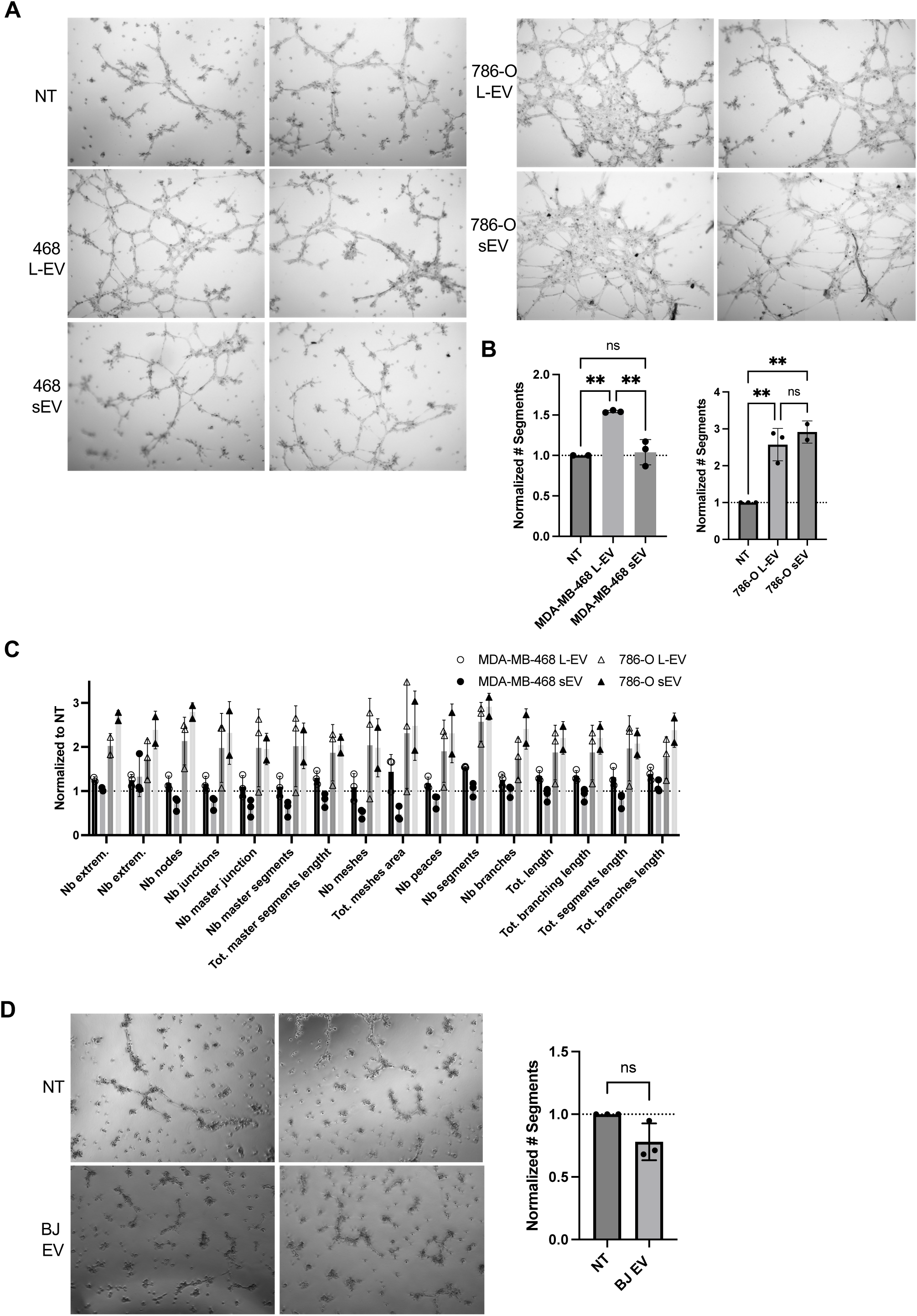
(A-C) L-EVs and sEVs were isolated from MDA-MB-468 and 786-O tumor cells, as described in the methods, and used to perform a tube assay. Networks were imaged after 5 hours and analyzed in Fiji with normalization to the no treatment control. The p-values were obtained by one-way ANOVA with Tukey’s correction. (D) EVs were isolated from non-tumorigenic BJ fibroblast cells and incubated with SVEC4-10 endothelial cells in a tube assay. Resulting networks were imaged after 5 hours, and number of segments was plotted normalized to the no treatment control as means ± SD. The p-value was obtained by an unpaired two-tailed t test. (ns: not significant, *p<0.05, ** p<0.01, ***p<0.001, ****p<0.0001). Bar graphs were constructed from at least 3 independent experiments.

**Supplemental Figure 4.**
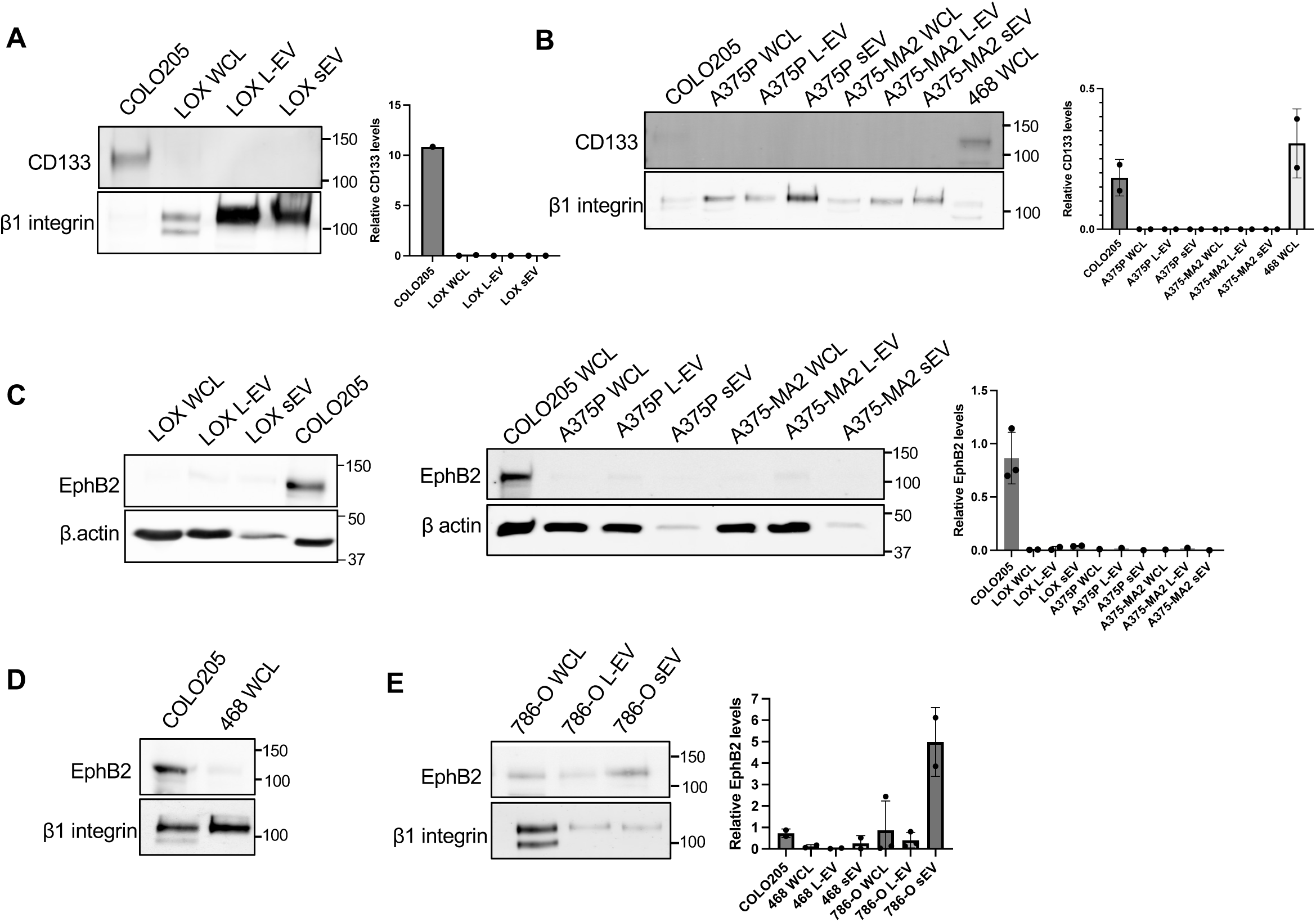
Corresponding whole cells (WCL), L-EVs and sEVs for each tumor cell line were lysed and separated by SDS-PAGE. The cargo contents were analyzed by western blotting. CD133 (A,B) and EphB2 (C-E) levels were compared to positive control lysates and β actin or β1 integrin were used as loading controls. Pixel densities were measured in Fiji. Data are presented as means ± SD. Molecular weight markers (kD) are indicated.

**Supplemental Figure 5.**
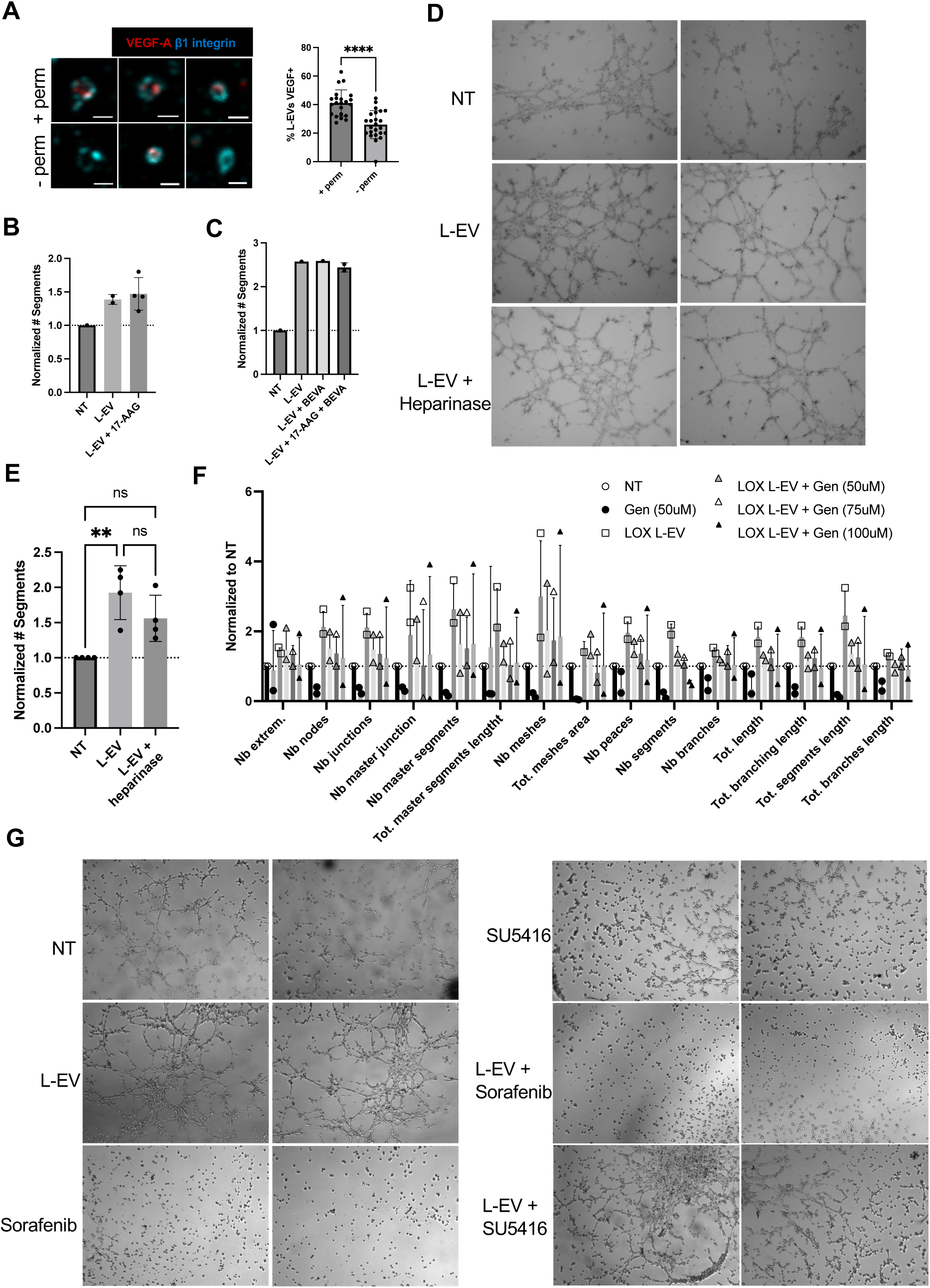
(A) L-EVs were purified from LOX melanoma cells as described in the methods and allowed to adhere on poly-_L_-lysine-coated coverslips. Vesicles were fixed and stained with VEGF-A and β1-integrin in the presence (+ perm) or absence (-perm) of Triton-X-100 and Tween-20. Scale bar, 0.5µm. Percent of vesicles that co-stained with VEGF-A under each condition was manually quantified and plotted as means ± SD. The experiment was repeated with quantitation of at least 22 fields per coverslip. The p-value was obtained by an unpaired two-tailed t test. (B) LOX cells were treated with 17-AAG (1μM) or DMSO control and allowed to shed for 24 hours. L-EVs were collected from the conditioned media by serial ultracentrifugation and used to perform a tube formation assay. Number of segments was quantified after 5 hours using Fiji and normalized to the no treatment condition. (C) L-EVs were pre-treated with 17-AAG (10μM) for 90 minutes at 37°C and then used in combination with bevacizumab (BEVA) (0.5μg/mL) for a tube assay. Number of segments was quantified using Fiji and normalized to the no treatment condition. (D,E) LOX L-EVs were pre-treated with heparinase (0.9mU/mL) for 16 hours prior to use with a tube formation assay. Number of segments was quantified after 5 hours using Fiji. The p-values were obtained by one-way ANOVA with Tukey’s correction. (F) Full tube network analysis for genistein (Gen) tube assay experiments. (G) SVEC4-10 cells were incubated with 10μM SU5416 or 20μM Sorafenib alone or in combination with L-EVs, cells allowed to form tubes and imaged after 5 hours. Independent biological replicates are presented as means ± SD. (ns: not significant, *p<0.05, ** p<0.01, ***p<0.001, ****p<0.0001).

**Supplemental Figure 6.**
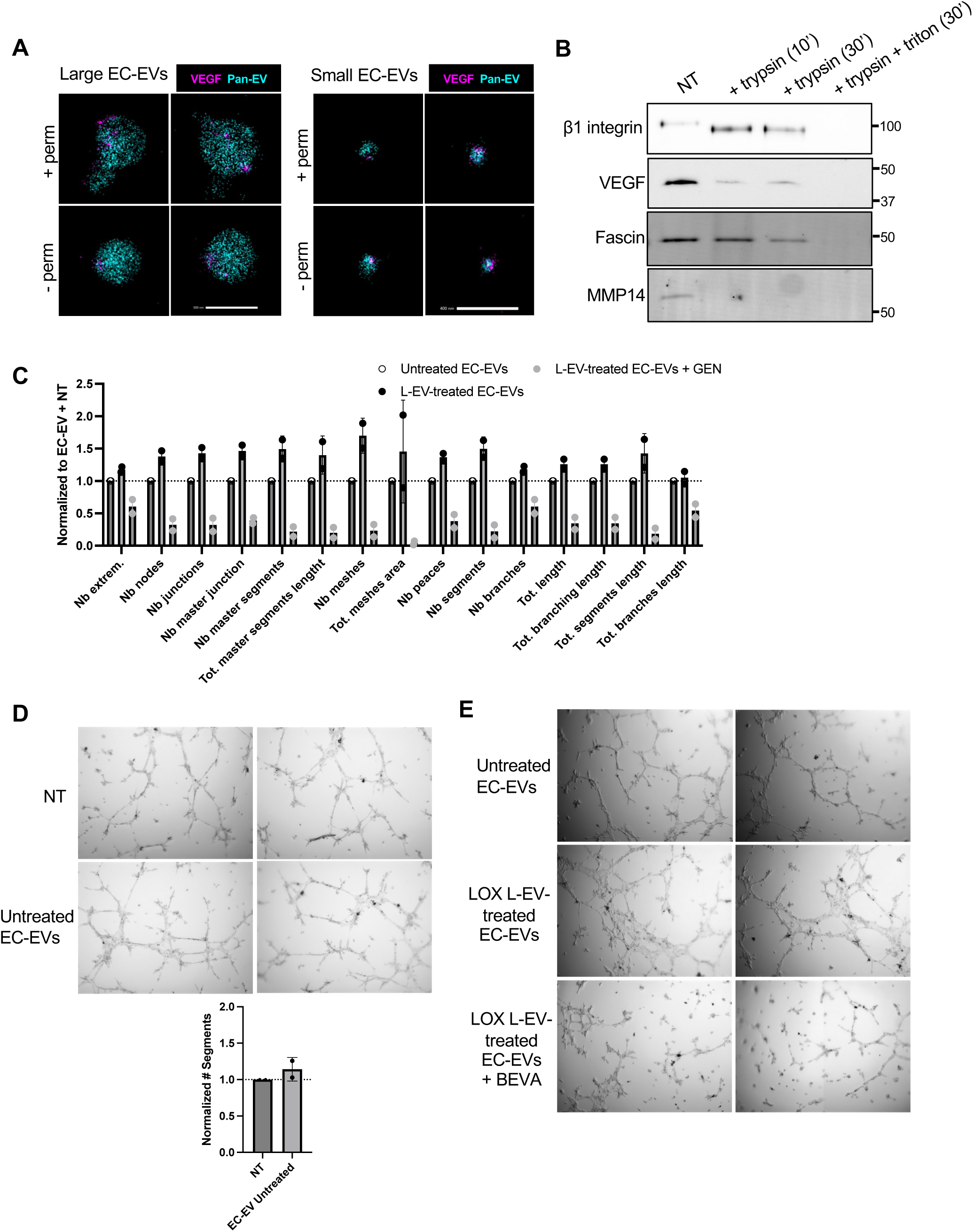
(A) EC-EVs were collected from LOX L-EV-treated HUVEC cells as described in the methods and processed for super resolution microscopy. Vesicles were fixed and stained with VEGF-A and Pan-EV in the presence (+ perm) or absence (-perm) of saponin. EC-EVs were binned as large (>200nm) or small (<200nm). Scale bars, 500nm (left) and 400nm (right). (B) EC-EVs were collected by ultracentrifugation and used to perform a trypsin digestion, as outlined in the methods. EC-EVs were re-isolated, re-suspended in 1.5X loading dye, and separated by SDS-PAGE for western blot analysis. Molecular weight markers (kD) are indicated. (C) Full tube network analysis for EC-EV experiments. (D) EC-EVs were collected from untreated HUVEC cells and used in a tube formation assay. Networks were imaged after 5 hours and number of segments was quantified and plotted as means ± SD after normalization. (E) EC-EVs were used alone or in combination with 100nM bevacizumab for a tube formation assay. Networks were imaged after 5 hours.

**Supplemental Figure 7.**
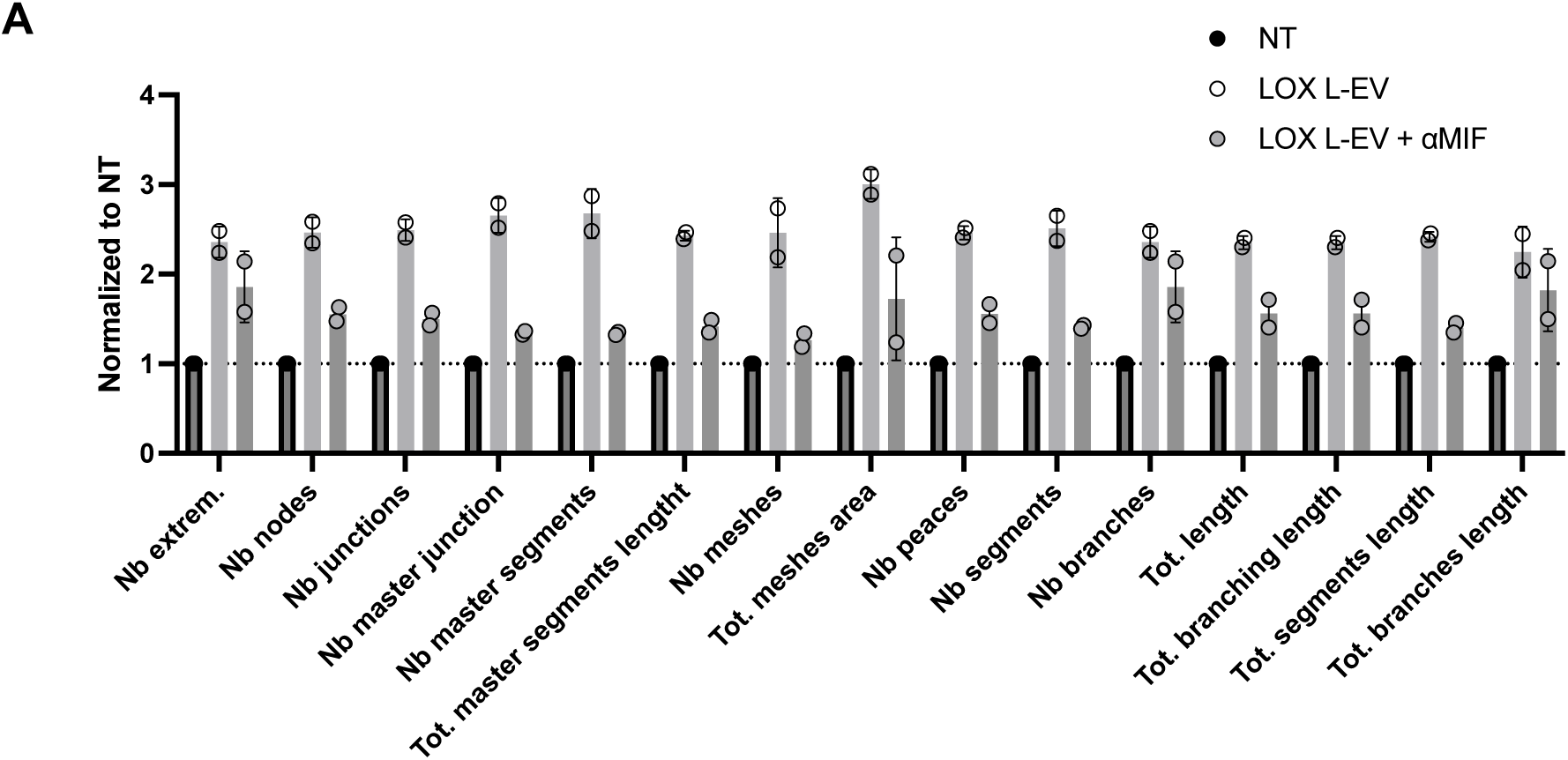
(A) Full tube network analysis for MIF neutralization experiments. Independent replicates are plotted as means ± SD after normalization.

## Notes

### Competing Interest Statement

The authors have declared no competing interest.

## REFERENCES

1. Waseh, S., and Lee, J. B. (2023) Advances in melanoma: epidemiology, diagnosis, and prognosis. Front Med (Lausanne). 10.3389/fmed.2023.1268479

2. Arnold, M., Singh, D., Laversanne, M., Vignat, J., Vaccarella, S., Meheus, F., Cust, A. E., De Vries, E., Whiteman, D. C., and Bray, F. (2022) Global Burden of Cutaneous Melanoma in 2020 and Projections to 2040. JAMA Dermatol. 158, 495–503

3. Arnold, M., de Vries, E., Whiteman, D. C., Jemal, A., Bray, F., Parkin, D. M., and Soerjomataram, I. (2018) Global burden of cutaneous melanoma attributable to ultraviolet radiation in 2012. Int J Cancer. 143, 1305–1314

4. Kashani-Sabet, M., Sagebiel, R. W., Ferreira, C. M. M., Nosrati, M., and Miller, J. R. (2002) Tumor vascularity in the prognostic assessment of primary cutaneous melanoma. Journal of Clinical Oncology. 20, 1826–1831

5. Ribatti, D., Annese, T., and Longo, V. (2010) Angiogenesis and Melanoma. Cancers (Basel). 2, 114–132

6. Liu, Z. L., Chen, H. H., Zheng, L. L., Sun, L. P., and Shi, L. (2023) Angiogenic signaling pathways and anti-angiogenic therapy for cancer. Signal Transduct Target Ther. 10.1038/s41392-023-01460-1

7. Lázár-Molnár, E., Hegyesi, H., Tóth, S., and Falus, A. (2000) Autocrine and paracrine regulation by cytokines and growth factors in melanoma. Cytokine. 12, 547–554

8. Latifkar, A., Hur, Y. H., Sanchez, J. C., Cerione, R. A., and Antonyak, M. A. (2019) New insights into extracellular vesicle biogenesis and function. J Cell Sci. 10.1242/JCS.222406

9. O’Brien, K., Breyne, K., Ughetto, S., Laurent, L. C., and Breakefield, X. O. (2020) RNA delivery by extracellular vesicles in mammalian cells and its applications. Nat Rev Mol Cell Biol. 21, 585–606

10. Minciacchi, V. R., Freeman, M. R., and Di Vizio, D. (2015) Extracellular Vesicles in Cancer: Exosomes, Microvesicles and the Emerging Role of Large Oncosomes. Semin Cell Dev Biol. 40, 41–51

11. Ko, S. Y., and Naora, H. (2020) Extracellular vesicle membrane-associated proteins: Emerging roles in tumor angiogenesis and anti-angiogenesis therapy resistance. Int J Mol Sci. 21, 1–17

12. Zhang, D. X., Vu, L. T., Ismail, N. N., Le, M. T. N., and Grimson, A. (2021) Landscape of extracellular vesicles in the tumour microenvironment: Interactions with stromal cells and with non-cell components, and impacts on metabolic reprogramming, horizontal transfer of neoplastic traits, and the emergence of therapeutic resistance. Semin Cancer Biol. 74, 24–44

13. Sheehan, C., and D’Souza-Schorey, C. (2019) Tumor-derived extracellular vesicles: Molecular parcels that enable regulation of the immune response in cancer. J Cell Sci. 10.1242/jcs.235085

14. Peng, Z., Tong, Z., Ren, Z., Ye, M., and Hu, K. (2023) Cancer-associated fibroblasts and its derived exosomes: a new perspective for reshaping the tumor microenvironment. Molecular Medicine. 10.1186/s10020-023-00665-y

15. Jeppesen, D. K., Zhang, Q., Franklin, J. L., and Coffey, R. J. (2023) Extracellular vesicles and nanoparticles: emerging complexities. Trends Cell Biol. 33, 667–681

16. Clancy, J. W., Boomgarden, A. C., and D’Souza-Schorey, C. (2021) Profiling and promise of supermeres. Nat Cell Biol. 23, 1216–1223

17. Bergers, G., and Hanahan, D. (2008) Modes of resistance to anti-angiogenic therapy. Nat Rev Cancer. 8, 592–603

18. Ye, Z. W., Yu, Z. L., Chen, G., and Jia, J. (2023) Extracellular vesicles in tumor angiogenesis and resistance to anti-angiogenic therapy. Cancer Sci. 114, 2739–2749

19. Zhou, J., Liu, X., Dong, Q., Li, J., Niu, W., and Liu, T. (2024) Extracellular vesicle-bound VEGF in oral squamous cell carcinoma and its role in resistance to Bevacizumab Therapy. Cancer Cell Int. 10.1186/s12935-024-03476-1

20. Ko, S. Y., Lee, W. J., Kenny, H. A., Dang, L. H., Ellis, L. M., Jonasch, E., Lengyel, E., and Naora, H. (2019) Cancer-derived small extracellular vesicles promote angiogenesis by heparin-bound, bevacizumab-insensitive VEGF, independent of vesicle uptake. Commun Biol. 10.1038/s42003-019-0609-x

21. Clancy, J. W., Sheehan, C. S., Boomgarden, A. C., and D’Souza-Schorey, C. (2022) Recruitment of DNA to tumor-derived microvesicles. Cell Rep. 10.1016/j.celrep.2022.110443

22. Montesano, R., Orci, L., and Vassalli, P. (1983) In Vitro Rapid Organization of Endothelial Cells into Capillary-like Networks Is Promoted by Collagen Matrices. J Cell Biol. 97, 1648–1652

23. Goodwin, A. M. (2007) In vitro assays of angiogenesis for assessment of angiogenic and anti-angiogenic agents. Microvasc Res. 74, 172–183

24. Carpentier, G., Berndt, S., Ferratge, S., Rasband, W., Cuendet, M., Uzan, G., and Albanese, P. (2020) Angiogenesis Analyzer for ImageJ — A comparative morphometric analysis of “Endothelial Tube Formation Assay” and “Fibrin Bead Assay.” Sci Rep. 10.1038/s41598-020-67289-8

25. Jeppesen, D. K., Fenix, A. M., Franklin, J. L., Higginbotham, J. N., Zhang, Q., Zimmerman, L. J., Liebler, D. C., Ping, J., Liu, Q., Evans, R., Fissell, W. H., Patton, J. G., Rome, L. H., Burnette, D. T., and Coffey, R. J. (2019) Reassessment of Exosome Composition. Cell. 177, 428–445.e18

26. Sato, S., Vasaikar, S., Eskaros, A., Kim, Y., Lewis, J. S., Zhang, B., Zijlstra, A., and Weaver, A. M. (2019) EPHB2 carried on small extracellular vesicles induces tumor angiogenesis via activation of ephrin reverse signaling. JCI Insight. 10.1172/jci.insight.132447

27. Feng, Q., Zhang, C., Lum, D., Druso, J. E., Blank, B., Wilson, K. F., Welm, A., Antonyak, M. A., and Cerione, R. A. (2017) A class of extracellular vesicles from breast cancer cells activates VEGF receptors and tumour angiogenesis. Nat Commun. 10.1038/ncomms14450

28. Gerhardt, H., Golding, M., Fruttiger, M., Ruhrberg, C., Lundkvist, A., Abramsson, A., Jeltsch, M., Mitchell, C., Alitalo, K., Shima, D., and Betsholtz, C. (2003) VEGF guides angiogenic sprouting utilizing endothelial tip cell filopodia. Journal of Cell Biology. 161, 1163–1177

29. Qian, C. N., Huang, D., Wondergem, B., and Teh, B. T. (2009) Complexity of tumor vasculature in clear cell renal cell carcinoma. in Cancer, pp. 2282–2289, 115, 2282–2289

30. Kim, B., Kim, S., Park, S., and Ko, J. (2024) CD133-containing microvesicles promote colorectal cancer progression by inducing tumor angiogenesis. Heliyon. 10.1016/j.heliyon.2024.e29292

31. Varker, K. A., Biber, J. E., Kefauver, C., Jensen, R., Lehman, A., Young, D., Wu, H., Lesinski, G. B., Kendra, K., Chen, H. X., Walker, M. J., and Carson, W. E. (2007) A randomized phase 2 trial of bevacizumab with or without daily low-dose interferon alfa-2b in metastatic malignant melanoma. Ann Surg Oncol. 14, 2367–2376

32. Sharifi-Rad, J., Quispe, C., Imran, M., Rauf, A., Nadeem, M., Gondal, T. A., Ahmad, B., Atif, M., Mubarak, M. S., Sytar, O., Zhilina, O. M., Garsiya, E. R., Smeriglio, A., Trombetta, D., Pons, D. G., Martorell, M., Cardoso, S. M., Razis, A. F. A., Sunusi, U., Kamal, R. M., Rotariu, L. S., Butnariu, M., Docea, A. O., and Calina, D. (2021) Genistein: An Integrative Overview of Its Mode of Action, Pharmacological Properties, and Health Benefits. Oxid Med Cell Longev. 10.1155/2021/3268136

33. Wilhelm, S. M., Adnane, L., Newell, P., Villanueva, A., Llovet, J. M., and Lynch, M. (2008) Preclinical overview of sorafenib, a multikinase inhibitor that targets both Raf and VEGF and PDGF receptor tyrosine kinase signaling. Mol Cancer Ther. 7, 3129–3140

34. Geindreau, M., Bruchard, M., and Vegran, F. (2022) Role of Cytokines and Chemokines in Angiogenesis in a Tumor Context. Cancers (Basel). 10.3390/cancers14102446

35. Li, A., Dubey, S., Varney, M. L., Dave, B. J., and Singh, R. K. (2003) IL-8 Directly Enhanced Endothelial Cell Survival, Proliferation, and Matrix Metalloproteinases Production and Regulated Angiogenesis. The Journal of Immunology. 170, 3369–3376

36. Wei, L.-H., Kuo, M.-L., Chen, C.-A., Chou, C.-H., Lai, K.-B., Lee, C.-N., and Hsieh, C.-Y. (2003) Interleukin-6 promotes cervical tumor growth by VEGF-dependent angiogenesis via a STAT3 pathway. Oncogene. 22, 1517–1527

37. Strozyk, E. A., Desch, A., Poeppelmann, B., Magnolo, N., Wegener, J., Huck, V., and Schneider, S. W. (2014) Melanoma-derived IL-1 converts vascular endothelium to a proinflammatory and procoagulatory phenotype via NFκB activation. Exp Dermatol. 23, 670–676

38. Clancy, J. W., and D’souza-Schorey, C. (2023) Tumor-Derived Extracellular Vesicles: Multifunctional Entities in the Tumor Microenvironment. Annu. Rev. Pathol. Mech. Dis. 18, 205–229

39. Chang, W. H., Cerione, R. A., and Antonyak, M. A. (2021) Extracellular Vesicles and Their Roles in Cancer Progression. in Methods in Molecular Biology, pp. 143–170, Humana Press Inc., 2174, 143–170

40. Huang, M., Lei, Y., Zhong, Y., Chung, C., Wang, M., Hu, M., and Deng, L. (2021) New Insights Into the Regulatory Roles of Extracellular Vesicles in Tumor Angiogenesis and Their Clinical Implications. Front Cell Dev Biol. 10.3389/fcell.2021.791882

41. Todorova, D., Simoncini, S., Lacroix, R., Sabatier, F., and Dignat-George, F. (2017) Extracellular vesicles in angiogenesis. Circ Res. 120, 1658–1673

42. Cui, C., Yan, X., Liu, S., Deitz, A. C., Si, L., Chi, Z., Sheng, X., Lian, B., Li, J., Ge, J., Wang, X., Mao, L., Tang, B., Zhou, L., Bai, X., Li, S., Li, B., Wu, H., and Guo, J. (2019) Real-world clinical outcomes of anticancer treatments in patients with advanced melanoma in China: retrospective, observational study. Int J Surg Oncol (N Y). 4, e76–e76

43. Han, X., Ge, P., Liu, S., Yang, D., Zhang, J., Wang, X., and Liang, W. (2023) Efficacy and safety of bevacizumab in patients with malignant melanoma: a systematic review and PRISMA-compliant meta-analysis of randomized controlled trials and non-comparative clinical studies. Front Pharmacol. 10.3389/fphar.2023.1163805

44. D’Souza-Schorey, C., and Di Vizio, D. (2025) A class of large cell-like extracellular vesicles: Extracellular vesicles. Nat Cell Biol. 27, 372–374

45. Atkin-Smith, G. K., Santavanond, J. P., Light, A., Rimes, J. S., Samson, A. L., Er, J., Liu, J., Johnson, D. N., Le Page, M., Rajasekhar, P., Yip, R. K. H., Geoghegan, N. D., Rogers, K. L., Chang, C., Bryant, V. L., Margetts, M., Keightley, M. C., Kilpatrick, T. J., Binder, M. D., Tran, S., Lee, E. F., Fairlie, W. D., Ozkocak, D. C., Wei, A. H., Hawkins, E. D., and Poon, I. K. H. (2024) In situ visualization of endothelial cell-derived extracellular vesicle formation in steady state and malignant conditions. Nature Communications. 10.1038/s41467-024-52867-5

46. Chesney, J., Metz, C., Bacher, M., Peng, T., Meinhardt, A., and Bucalal, R. (1999) An Essential Role for Macrophage Migration Inhibitory Factor (MIF) in Angiogenesis and the Growth of a Murine Lymphoma The role of MIF in. Molecular Medicine. 5, 181–191

47. Mitchell, R. A., and Bucala, R. (2000) Tumor growth-promoting properties of macrophage migration inhibitory factor (MIF). Semin Cancer Biol. 10, 359–366

48. Shi, J., and Wei, P. K. (2016) Interleukin-8: A potent promoter of angiogenesis in gastric cancer. Oncol Lett. 11, 1043–1050

49. Ugurel, S., Rappl, G., Tilgen, W., and Reinhold, U. (2001) Increased Serum Concentration of Angiogenic Factors in Malignant Melanoma Patients Correlates With Tumor Progression and Survival. Journal of Clinical Oncology. 19, 577583

50. Kargl, J., Zhu, X., Zhang, H., Yang, G. H. Y., Friesen, T. J., Shipley, M., Maeda, D. Y., Zebala, J. A., McKay-Fleisch, J., Meredith, G., Mashadi-Hossein, A., Baik, C., Pierce, R. H., Redman, M. W., Thompson, J. C., Albelda, S. M., Bolouri, H., and McGarry Houghton, A. (2019) Neutrophil content predicts lymphocyte depletion and anti-PD1 treatment failure in NSCLC. JCI Insight. 10.1172/jci.insight.130850

51. Bilusic, M., Heery, C. R., Collins, J. M., Donahue, R. N., Palena, C., Madan, R. A., Karzai, F., Marté, J. L., Strauss, J., Gatti-Mays, M. E., Schlom, J., and Gulley, J. L. (2019) Phase i trial of HuMax-IL8 (BMS-986253), an anti-IL-8 monoclonal antibody, in patients with metastatic or unresectable solid tumors. J Immunother Cancer. 10.1186/s40425-019-0706-x

52. Ismail, A. A., Shaker, B. T., and Bajou, K. (2022) The plasminogen–activator plasmin system in physiological and pathophysiological angiogenesis. Int J Mol Sci. 10.3390/ijms23010337

53. Wang, X., Montoyo-Pujol, Y. G., Bermudez, S., Corpas, G., Martin, A., Almazan, F., Cabrera, T., and López-Nevot, M. A. (2021) Serum Cytokine Profiles of Melanoma Patients and Their Association with Tumor Progression and Metastasis. J Oncol. 10.1155/2021/6610769

54. Laino, A. S., Woods, D., Vassallo, M., Qian, X., Tang, H., Wind-Rotolo, M., and Weber, J. (2020) Serum interleukin-6 and C-reactive protein are associated with survival in melanoma patients receiving immune checkpoint inhibition. J Immunother Cancer. 10.1136/jitc-2020-000842

